# The Parkinson’s drug entacapone disrupts gut microbiome homeostasis via iron sequestration

**DOI:** 10.1101/2023.11.12.566429

**Authors:** Fátima C. Pereira, Xiaowei Ge, Jannie Munk Kristensen, Rasmus H. Kirkegaard, Klara Maritsch, Yifan Zhu, Marie Decorte, Bela Hausmann, David Berry, Kenneth Wasmund, Arno Schintlmeister, Thomas Boettcher, Ji-Xin Cheng, Michael Wagner

## Abstract

Increasing evidence shows that many human-targeted drugs alter the gut microbiome, leading to implications for host health. However, much less is known about the mechanisms by which drugs target the microbiome and how drugs affect microbial function. Here we combined quantitative microbiome profiling, long-read metagenomics, stable isotope probing and single cell chemical imaging to investigate the impact of two widely prescribed nervous system targeted drugs on the gut microbiome. *Ex vivo* supplementation of physiologically relevant concentrations of entacapone or loxapine succinate to faecal samples significantly impacted the abundance of up to one third of the microbial species present. Importantly, we demonstrate that the impact of these drugs on microbial metabolism is much more pronounced than their impact on abundances, with low concentrations of drugs reducing the activity, but not the abundance of key microbiome members like *Bacteroides, Ruminococcus* or *Clostridium* species. We further demonstrate that entacapone impacts the microbiome due to its ability to complex and deplete available iron, and that microbial growth can be rescued by replenishing levels of microbiota-accessible iron. Remarkably, entacapone-induced iron starvation selected for iron-scavenging organisms carrying antimicrobial resistance and virulence genes. Collectively, our study unveils the impact of two under-investigated drugs on whole microbiomes and identifies metal sequestration as a mechanism of drug-induced microbiome disturbance.

## Main

Drugs initially designed to specifically target human cells often can affect microbes as well^1^. As a result of poor gastrointestinal absorption and/or biliary secretion, many of these drugs reach the large intestine where they encounter - and potentially interact with - hundreds to thousands of different microbial species that play important roles in various aspects of human physiology^1,2,3^. Indeed, several cohort studies have reported significant associations between the use of medication and shifts in gut microbial composition and function^4,5,6,7^. While the cross-sectional nature of these large cohort studies hinders the establishment of causality, *in vitro* studies have been pivotal for systematically evaluating direct effects of human-targeted drugs on gut microbes. A landmark study assessing the antimicrobial effect of 835 human-targeted drugs against a panel of 40 cultured gut microbes revealed that a significant proportion (24%) of drugs could inhibit the growth of at least one gut bacterial strain *in vitro*^1^. An additional study testing the effect of a smaller panel of drugs on faecal samples by metaproteomics demonstrated selective anti- and/or pro-microbial activity for the great majority of drugs tested, with a significant fraction of drugs also shifting microbiome composition^8^. Of note, this study demonstrated that bacterial function could shift in response to drugs without a change in taxon abundance, thus highlighting the need for using metrics other than abundance when investigating the impact of drugs. Importantly, the interaction between the microbiome and drugs is bidirectional, with many studies clearly demonstrating that gut microbes can also actively metabolize^9,10,11,12^, and under certain circumstances bioaccumulate^13^, pharmaceutical drugs. While several studies to date have shed light on the nature and extent of microbe-driven drug transformations^9,10,11^, mechanistic details for human-targeted drug impact on the microbiome remain to be elucidated.

One hypothesis is that drugs can change intestinal microenvironments such as pH or osmolarity, and by doing so, directly affect bacterial growth^14^. Another explanation is that drugs interact with structural analogues of their human targets within bacteria, thus interfering with cellular processes also in microbes^15^. Drug-microbiome interactions have been shown to modulate the therapeutic effect of the drug, contribute to its side effects, or both^11,16,17,18^. Importantly, in certain cases, drug-induced microbiome changes might also contribute to other diseases. For instance, proton pump inhibitors (PPIs) cause major shifts in the gut microbiome^19^, leading to decreased resistance to colonisation by enteric pathogens such as *Clostridioides difficile*, *Campylobacter* and *Salmonella*^20^. Furthermore, PPIs-induced shifts in the microbiome during early childhood have been linked to obesity^21^.

Among the human-targeted drugs tested *in vitro*, compounds that target the nervous system seem to exhibit stronger anti-commensal activity against gut bacteria compared to other tested drug classes^1^. Indeed, several studies have reported a strong and selective inhibitory activity of antipsychotics and antidepressants on gut microbial strains and microbiomes^22,23,24^. This is concerning, given the growing number of studies implicating the microbiome in many neuropsychiatric disorders^25^, and the widespread and rising use of this class of pharmaceuticals worldwide^26^. Thus, a better mechanistic understanding of drug-microbiome interactions in the context of nervous system-targeted medications may facilitate novel ways to improve efficacy and/or minimize side effects of therapies for such disorders.

Here, we investigate the effects of two nervous system-targeted drugs on whole gut microbiomes using a suite of complementary functional microbiome approaches. These aimed at investigating the effects of the drugs in the context of whole microbial communities, which more closely resembles their effects *in situ*, and to therefore examine most key members of the gut microbiome. We studied two commonly used drugs: i) entacapone, a catechol-O-methyltransferase (COMT) inhibitor that acts by preventing the degradation of levodopa, the main drug used in the treatment of Parkinson’s disease^27^; ii) loxapine succinate, a tricyclic antipsychotic medication primarily used in the treatment of schizophrenia^28^. These drugs, whose yearly total prescription exceeds 30 million tablets only in the US^29^, belong to different therapeutic classes and were shown to be selective in their anti-commensal activity against a panel of 40 cultured gut strains, with entacapone predominantly targeting Gram-positive organisms of the Firmicutes phylum and loxapine succinate only taxa within the Gram-negative order Bacteroidales^1^. We demonstrate that the impact of either drug on microbiome composition and/or activity extends beyond the taxa initially detected *in vitro* in pure culture, with entacapone causing strong shifts in microbiome composition. This prompted us to look for the cause of these shifts. We identified microbial iron deprivation, driven by the ability of entacapone to complex iron, as the main mechanism behind entacapone’s strong modulatory effect. These results advance our understanding of the impact of nervous-system targeted drugs on whole microbial communities and reveal micronutrient-deprivation as a mechanism through which entacapone disrupts the microbiome.

## Results

### Nervous-system targeted drugs affect microbiome composition and abundance

To evaluate the impact of entacapone (ENT) and loxapine succinate (LOX) on whole gut microbiome communities, we incubated freshly collected faecal samples from healthy adult individuals in sM9 medium with two different concentrations of each drug dissolved in dimethylsulfoxide (DMSO) (Fig. 1a, see Methods). The low drug concentration (ENT-Low, LOX-Low, 20 μM) was previously used in a screening aimed at determining drug effects on pure culture isolates, while the high concentrations (ENT-Hi, 1965 μM, LOX-Hi, 100 μM) were based on estimated colon concentrations for each drug, and were included to better reflect the exposure of gut microbes to these drugs in the large intestine^1^ (see Methods). After short incubation times (6 and 24 hours) under anaerobic conditions at 37 °C, samples were collected and processed to determine: i) changes of the total microbial loads; ii) microbial community profile dynamics based on 16S rRNA gene amplicon sequencing; iii) reconstructed microbial genomes based on long-read metagenomics and iv) single-cell microbial activity changes via tracing of the incorporation of deuterium from isotopically-labelled heavy water (D2O) into single cells of microbiota by chemical imaging based on stimulated Raman scattering spectroscopy (SRS)^30^ (Fig. 1a).

**Figure 1.**
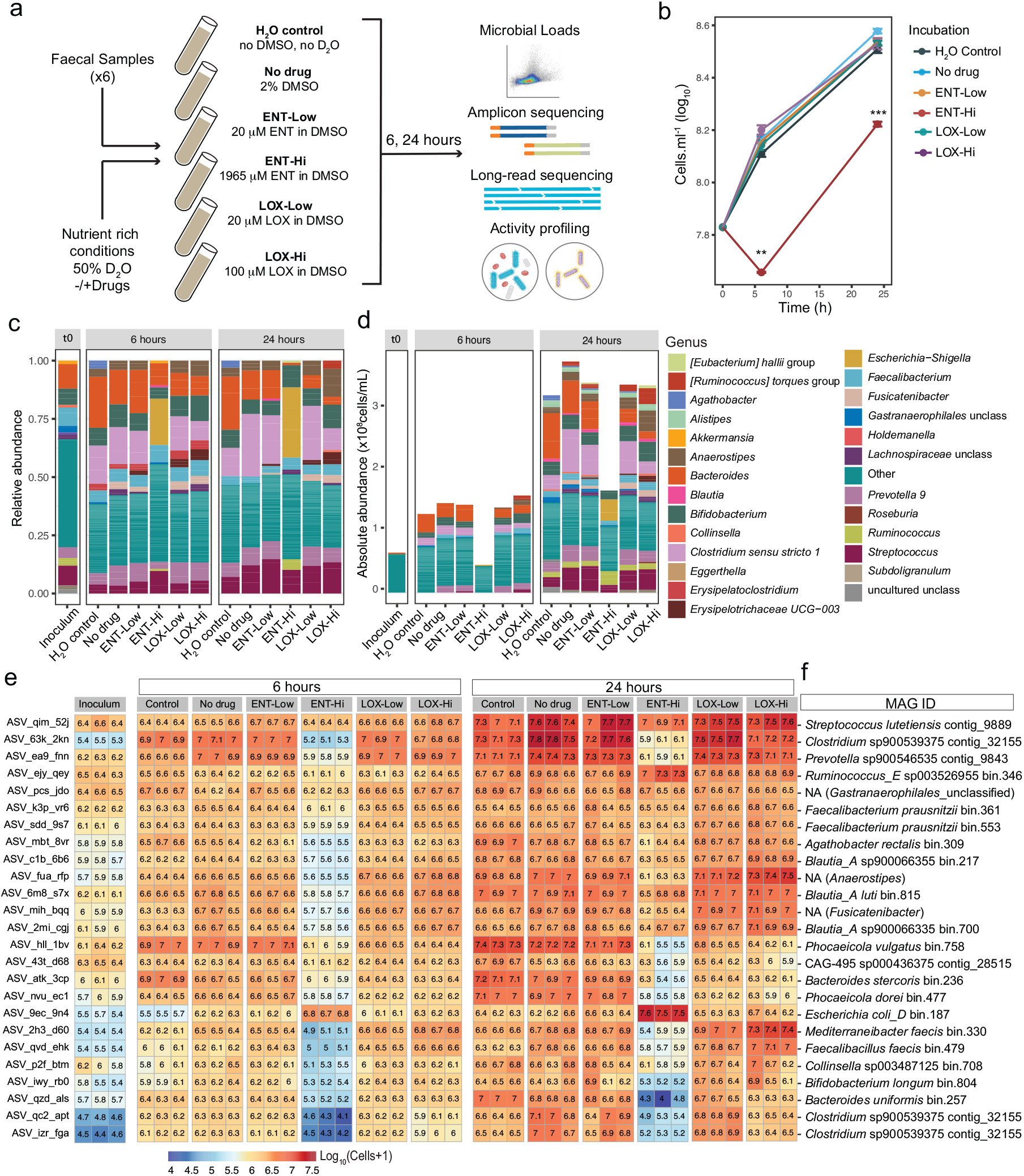
Drug-supplementation of faecal samples impacts biomass accumulation and microbiota composition. **a.** Schematic representation of faecal sample incubations with the drugs entacapone (ENT) and loxapine (LOX). As indicated, two different concentrations of each drug were used (Low and Hi). Incubations were performed in the presence of 50% heavy water (D_2_O) and 2% DMSO in the medium (either sM9 or BHI), except for the H_2_O control that consisted of medium without D_2_O or DMSO. After 6 or 24 hours of incubation, samples were collected for further analyses (see Methods). All incubations were performed in triplicates. **b.** Total microbial cell loads in faecal sample incubations described in (A), at the start of the incubation (time=0 hours) and after 6 and 24 hours, respectively. Total cell loads were assessed by flow cytometry. **p<0.01; ***p<0.001; unpaired two-sample t-test with “No drug” used as the reference. **c, d.** Relative (**c**) and absolute (**d**) genus abundance profiles of faecal microbial communities incubated for 6 or 24 hours as described in (**a**) and assessed by 16S rRNA gene amplicon sequencing. The community composition at 0 hours (inoculum) is also shown. All genera present at a relative abundance below 0.025 or absolute abundance below 6×10^6^ cells.ml^-1^ were assigned to the category “Other”. “unclass”: unclassified. **e**. Heatmap showing the absolute abundance, expressed in log_10_(cells+1), of the 25 most abundant ASVs detected across all samples. Each column displays data from one replicate. **f**. Metagenome-assembled genomes (MAGs) with 16S rRNA gene sequences matching ASVs shown in (**e**). MAGs were retrieved by metagenomic sequencing of the initial faecal samples using Oxford Nanopore sequencing (see Methods and Supplementary Table 7). ‘NA” indicates no match of the ASV sequence to each MAG 16S rRNA gene sequence.

Counts of total microbial cell loads by flow cytometry in samples incubated with or without drugs revealed that supplementation of a physiologically relevant concentration of entacapone (ENT-Hi) significantly reduced the numbers of microbial cells over time, when compared to all other tested conditions (Fig. 1b, Supplementary Table 1). Major shifts in the microbial community composition, as determined by 16S rRNA gene amplicon sequencing analyses, were detected in response to ENT-Hi, LOX-Hi and LOX-Low treatments (Fig. 1c), and in the case of ENT-Hi these were also accompanied by shifts in alpha diversity (Extended Data Fig. 1a,b; Supplementary Table 2,3).

By integrating total microbial counts with 16S rRNA gene amplicon sequencing data^31^, we determined absolute abundances for all detected taxa (Fig. 1d). Importantly, absolute abundance data confirmed that the employed incubation conditions enabled an increase in abundance for nearly all the taxa initially present in faecal samples (Fig. 1e: No drug versus inoculum; Supplementary Table 4) and thus allowed tracking drug-induced activity and abundance changes of microbiome members. Absolute abundances of genera such as *Bacteroides* or *Clostridium sensu stricto* 1 decreased in both ENT-Hi and LOX-Hi samples compared to the DMSO control (Fig. 1d). However, many of the detected effects were drug-specific, with ENT-Hi decreasing and LOX-Hi increasing total abundances of the genera *Anaerostipes, Fusicatenibacter*, *Ruminococcus torques* group, *Eubacterium hallii* group, *Erysipelotrichaceae* group UG-003 and *Roseburia*. Abundances of several genera were significantly altered in response to ENT-Hi only: *Escherichia-Shigella and Ruminococcus* increased in abundance, and genera such as *Alistipes*, *Streptococcus* or *Blautia* decreased (Fig. 1d,e; Supplementary Table 5). A similar impact of ENT-Hi on microbial biomass accumulation and on the overall community composition could also be observed when we used a different, nutrient rich medium (BHI) for the incubations (Extended Data Fig. 1c-g).

Differential abundance analysis indicated that the abundance of 29.4% of all 16S rRNA gene amplicon sequencing variants (ASVs) were significantly impacted by ENT-Hi after 24 hours of incubation, 11.8% of ASVs were impacted by LOX-Hi, and only 3.6% and 6.0% of ASVs were impacted by ENT-Low and LOX-Low, respectively (see Methods, Supplementary Table 5). Interestingly, LOX-Low resulted in growth inhibition patterns that differ from the ones observed when the same drug was supplemented to gut members grown under isolation^1^, where it only specifically inhibits growth of Bacteroidales strains. Our results show that other Gram-negatives and several Gram-positive species are affected by LOX-Low in the context of whole microbiome communities (Supplementary Table 5). These include *Erysipelotrichaceae* spp., *Oscillospiraceae* spp. and *Lachnospiraceae* spp., suggesting a cross-sensitization to loxapine in the context of the community. For ENT-Low, only a very low number of organisms were significantly impacted and were mostly Firmicutes, which agrees with previous reports^1^ (Supplementary Table 5).

Using long-read metagenomic sequencing of the starting faecal sample material, we retrieved a total of 1049 metagenome-assembled genomes (MAGs), 11 of which are complete genomes and 255 are medium-or high-quality genomes (Supplementary Tables 6 and 7). BLASTn^32^ analysis enabled us to link 16S rRNA gene sequences of ASVs to MAGs that contained 16S rRNA genes and to follow drug-driven community shifts at a higher taxonomic resolution (Fig. 1e,f). This revealed that ASVs classified as *Escherichia* and *Ruminococcus* taxa thriving in ENT-Hi give exact hits to the 16S rRNA genes in genomes of *E. coli* and *R. bromii*. These analyses also indicated that LOX-Hi conditions selectively promoted the growth of taxa like *Mediterraneibacter faecis*, *Faecalibacillus faecis* and *Blautia_A*. Species such as *Clostridium* sp900539375 (*Clostridium sensu stricto* 1 based on SILVA taxonomy) and several *Bacteroides* species *(B. uniformis, B. stercoris, Phocaeicola dorei* - formerly *B. dorei,* and *P. vulgatus* - formely *B. vulgatus*) were totally or partially inhibited by the presence of high concentrations of either drug, but the effect of ENT-Hi is much more pronounced than that of LOX-Hi (Fig. 1e,f). On the other hand, *Prevotella* sp900546535 and several Gram-positive organisms such as *Streptococcus lutetiensis*, *Collinsella* sp003487125, *Bifidobacterium longum*, *Mediterraneibacter faecis* or *Faecalibacillus faecis* showed growth inhibition by ENT-Hi only (Fig. 1e,f). All together these results reveal a strong but distinct impact of entacapone and loxapine at physiological concentrations on microbiota composition and abundance, with entacapone having a much more pronounced effect than loxapine.

### Nervous system-targeted drugs can alter microbial metabolism without impacting microbial abundance

Next, we evaluated the effect of these drugs on microbial activity at the single-cell level and explored whether drug exposure that did not cause major shifts in microbiome abundance within 24 hours of incubation did, nevertheless, impact their activity. To determine drug-induced changes in microbial activity, we added D2O as universal metabolic tracer^33,34,35^ to our incubations (Fig. 1a). In complex microbial communities, all metabolically active cells will incorporate deuterium (D) from D2O into their biomass during synthesis of new macromolecules^34^. The newly formed carbon-deuterium (C-D) bonds can then be used as a read-out of microbial activity. Detection and quantification of C-D levels in single microbial cells can be achieved using SRS, a method that efficiently excites the Raman active vibrational modes coherently with two synchronized ultrafast lasers^36,37,38^. We have successfully combined SRS with fluorescence *in situ* hybridization (FISH) in the past to determine gut microbiome response to sugars with high-throughput^30^. Using an optimized SRS-FISH platform that provides even higher throughput and sensitivity than the previous setup (Supplementary Information text, Extended Data Fig. 2), we measured around thirty thousand individual microbiota cells after 6 and 24 hours of incubation in the presence of the drugs and in controls (Fig. 2a,b). An unexpected, non-vibrational signal was detected in samples that were incubated with ENT-Hi and thus these samples were excluded from SRS-based activity measurements and the origin of this signal was further explored as detailed in the next section. As expected, in the absence of the drugs, nearly all of the analyzed cells were detected as active in the incubation medium, with 90% and 98% of cells displaying %CDSRS values above threshold at 6 hours and 24 hours of incubation, respectively (Fig. 2b, Supplementary Table 8). Addition of either ENT-Low, LOX-Low or LOX-Hi resulted in a significantly reduced fraction of total active cells, as well as in a significant reduction of single-cell metabolic activity. This reduction was more pronounced for LOX-Hi, followed by LOX-Low and ENT-Low (Fig. 2b). Thus, these drugs clearly impacted microbial activity within communities, even after short incubation times and under conditions where no impacts on their overall abundances were detected (Fig. 1b).

**Figure 2.**
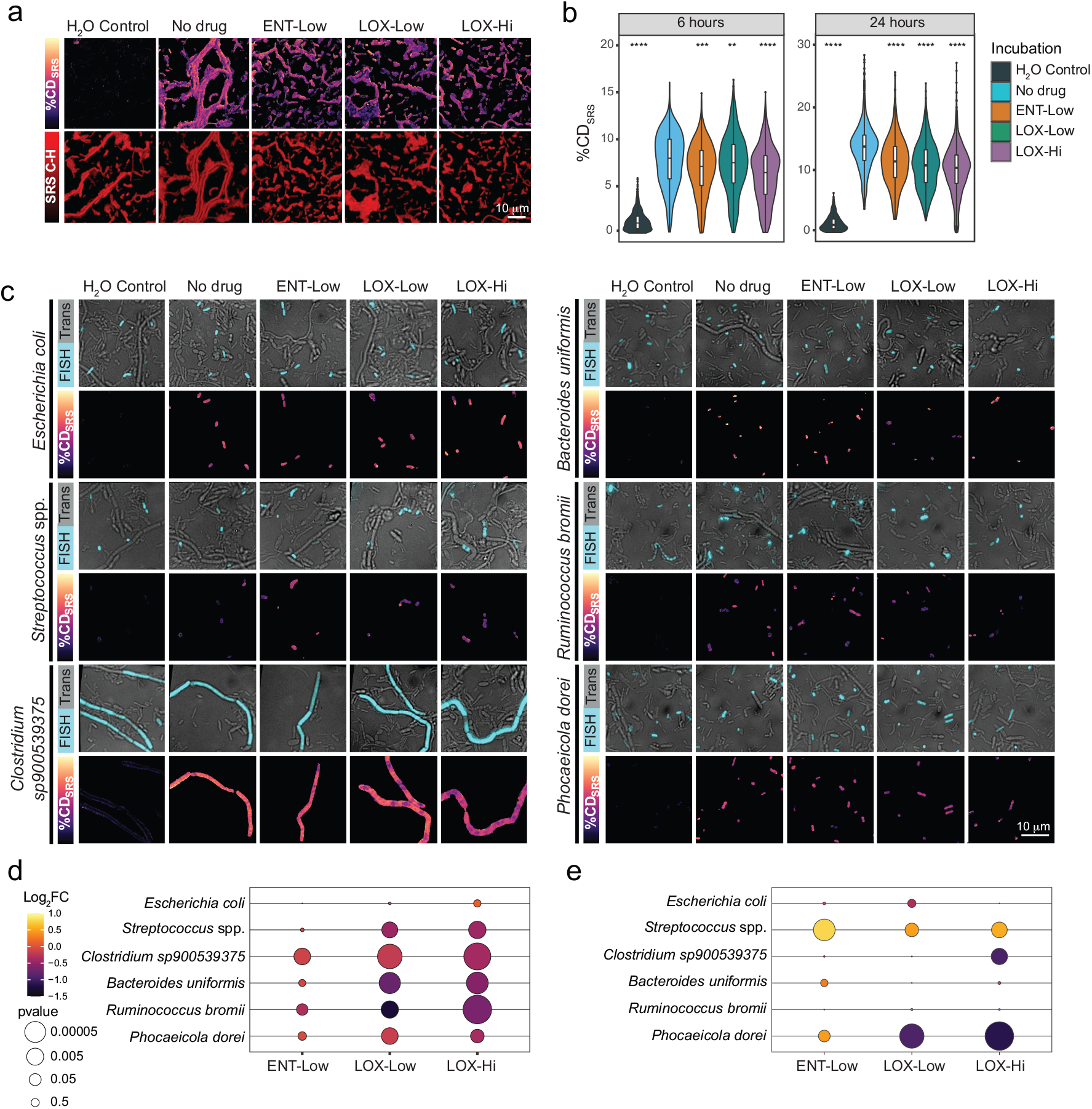
Metabolic activity of drug-incubated single-microbiome cells measured by deuterium incorporation via SRS. **a.** Representative SRS images of faecal samples incubated with the indicated drugs for 6 hours. The measure %CD_SRS_ indicates different levels of microbial activity. %CD_SRS_ distribution images are displayed on top and C-H for biomass visualization (log scaled) on the bottom. %CD_SRS_ scaling: minimum 0, maximum 20%. **b.** Single-cell %CD_SRS_ values in each analyzed sample. Violin plots illustrating summary statistics (median, first and third quartiles, with the extended lines representing the minimum and maximum values within 1.5 interquartile ranges from the first and third quartiles). **p=0.0043, ***p=5.2e-12, **** p<2e-16, Wilcoxon test using “No drug” as the reference. **c.** Drug-incubated faecal samples were hybridized with fluorescently-labeled oligonucleotide probes targeting *E. coli* (Ecol_268), *Streptococcus* and *Lactococcus* species (Strc_493), *Clostridium* sp900539375 (*Clostridium sensu stricto 1,* Clo_648), *Ruminococcus bromii* (Brom_2036), *B. uniformis* (Buni_1001), and *Phocaeicola dorei* (Bado_374) (all shown in cyan, Supplementary Table 9). For each targeted group, top rows show representative images obtained by overlay of transmitted light (grey) and fluorescence intensity (cyan). Bottom rows show the corresponding SRS images (displaying %CD_SRS_) for the FISH-targeted microbes (%CD_SRS_ values of other microbes are not displayed for the sake of visibility). %CD_SRS_ scaling: minimum 0, maximum 20%. Scale bar, 10 μm. **d.** Bubble plot denoting the fold change (FC, represented as Log_2_FC) in activity levels (calculated as %CD_SRS_) for the taxa targeted by FISH and incubated with drugs relative to “No drug” incubations. **e.** Bubble plot denoting the FC in absolute abundances for the taxa targeted by FISH and incubated with drugs relative to “No drug” incubations, as determined by DeSeq2.

To examine the activity of specific populations of the microbiome within the complex communities, we targeted six distinct abundant taxa of interest using SRS-FISH, which enabled us to determine the activity of individual cells from these taxa. Both published as well as newly designed and optimized oligonucleotide probes targeting 16S and 23S rRNA sequences predicted from MAGs retrieved by metagenomics were used for this purpose (Supplementary Table 9). Targeted taxa included organisms whose abundances were both negatively and positively impacted by drugs. We then evaluated the levels of activity of the targeted populations using picosecond SRS (Fig. 2c, Extended Data Fig. 2). Both LOX-Low and LOX-Hi incubations significantly reduced the activity of all targeted gut microbiota members except for *E. coli* (Fig. 2d). Interestingly, *Clostridium* sp900539375, *B. uniformis*, *Ruminococcus bromii* and *P. dorei* show reduced activity in LOX treatments, but only the abundance of *P. dorei* and *Clostridium* sp900539375 was negatively impacted (Fig. 2d,e; Supplementary Table 10). LOX-Low and LOX-Hi seem to strongly inhibit *P. dorei* growth within 6 hours (Fig. 1e), probably rendering most cells of this taxon undetectable by FISH, as the low ribosome content of non-active cells hinders FISH detection. We speculate that only a few drug-resistant *P. dorei* cells remained active enough to be detected by FISH, and these cells were not strongly impacted in activity (Fig. 2c,d). A comparable decrease in activity was detected for *Streptococcus* spp., but in this case this decrease was surprisingly accompanied by a slight increase in abundance at 6 hours (Fig. 2d,e). However, from 6 to 24 hours, we detect a decrease of the population represented by *Streptococcus lutentiensis* MAG in the presence of LOX compared to the control (Fig. 1e,f), which could be an effect of the lower activities detected by SRS in the LOX conditions at 6 hours. In summary, SRS-FISH provided new insights into the impact of ENT and LOX on the activities of specific microbiome members, which are often masked when only considering abundance data from relatively short incubation experiments.

### Entacapone bioaccumulates in microbiota cells

Gut microbes have been shown to bioaccumulate some drugs leading to depletion of the drug from the surrounding environment^13,39^. As ENT-Hi samples showed a strong, Raman unspecific signal during SRS pump-probe detection, we explored the origin of this signal and concluded that this is a photothermal (PT) signal originating from entacapone bioaccumulation within microbial cells (Supporting Information Text). By mapping the intensity of this PT signal from ENT-Hi samples and controls (Fig. 3a), we were able to show that the signal from entacapone occurred in a large fraction of microbial cells that had been incubated with the drug and washed before being fixed for analysis, but only in very few cells that had been fixed prior to incubation with the drug (i.e., dead cells) (Fig. 3b,c, Supplementary Table 11).

**Figure 3.**
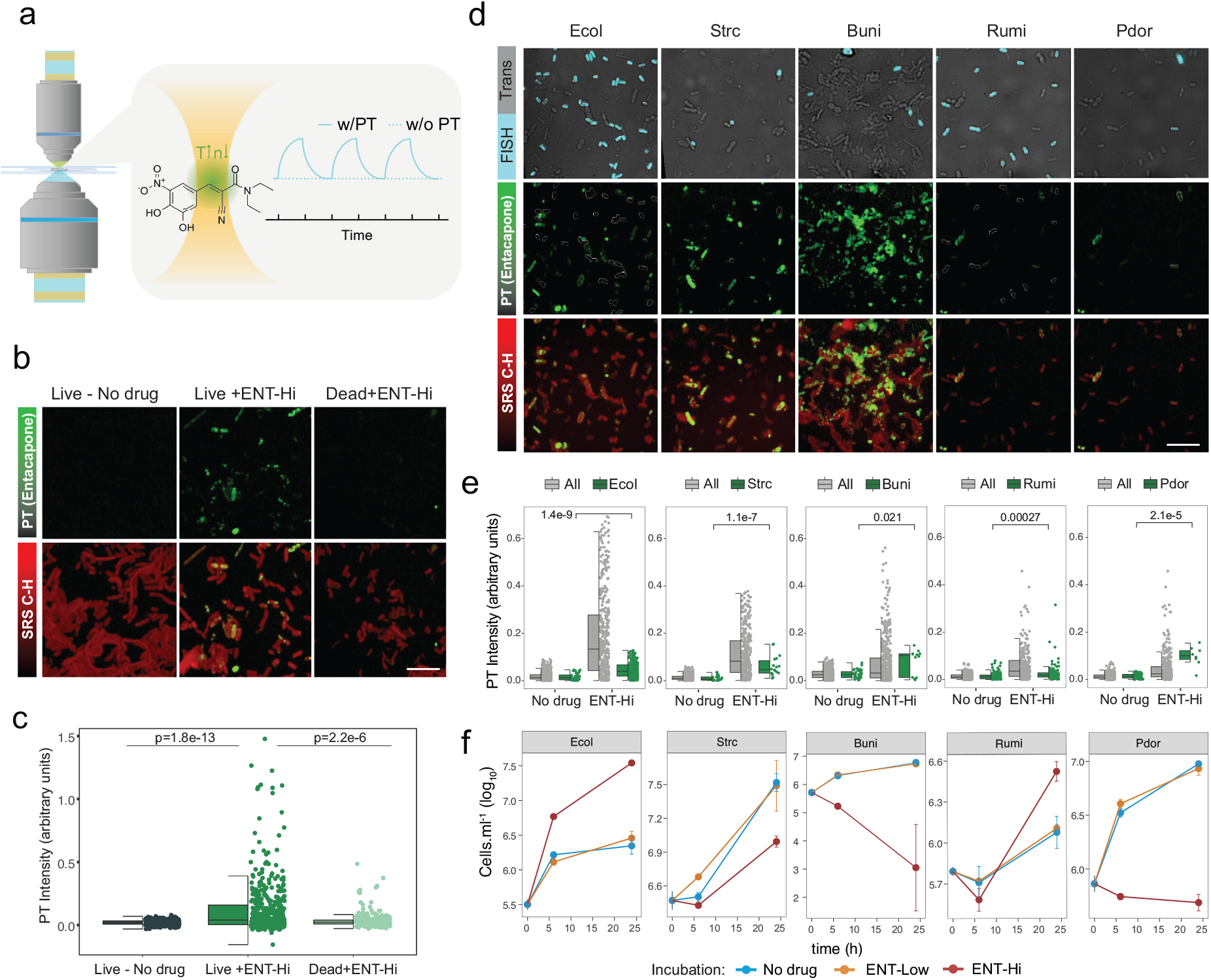
Photothermal imaging of entacapone bioaccumulation by microbiota cells. **a.** Schematic illustration of the time-dependent signal obtained from a solution of 10 mM entacapone in DMSO, with photothermal (w/PT) and without photothermal (w/o PT) detection. **b.** Photothermal signal intensity distribution from entacapone, shown in green (PT, determined by photothermal imaging), and C-H signal, shown in red, of live and dead (PFA fixed) microbiota cells incubated in the presence (+) or absence (-) of ENT-Hi for 6 hours. PT channel contrast: min 0 max 1.8. C-H signals are represented on a log scale. Scale bar, 10 μm. **c.** Single-cell photothermal signal intensity distribution in samples shown in (a). p-values were determined using an unpaired two-sample Wilcoxon test. **d.** Representative images of faecal samples incubated with entacapone for 6 hours followed by hybridization with fluorescently-labeled oligonucleotide probes targeting *E. coli* (Ecol), *Streptococcus* and *Lactococcus* (Strc), *Ruminococcus bromii* (Rumi), *B. uniformis* (Buni), and *Phocaeicola dorei* (Pdor) (see Supplementary Table 9). FISH-probe signals from hybridized cells (in cyan) were detected using widefield fluorescence microscopy (top panel). Photothermal signal maps from entacapone (PT, in green) are shown in the middle (cells targeted by FISH are shown with respective white contour lines). Entacapone PT signals overlayed with SRS C-H signals (in red) are shown in the bottom panel. PT channel contrast: min 0 max 1.8. C-H signals are represented on a log scale. Scale bar, 8 μm. **e.** Single-cell photothermal signal distribution in samples shown in (c). p-values were determined using the unpaired two-sample Wilcoxon test. **f.** Time series of the absolute abundance of the respective taxa targeted by FISH in No drug, Ent-Low and ENT-Hi incubations. In **c**, **e** boxes represent the median, first and third quartile. Whiskers extend to the highest and lowest values that are within one and a half times the interquartile range.

To identify the main drug accumulating taxa, we further looked at the entacapone distribution among populations targeted with the FISH probes described above (Fig. 3d, Supplementary Table 9). All of the targeted populations displayed entacapone signals to different intensities. While *P. dorei* and, to a lesser extent *Streptococcus* spp. and *E. coli,* seemed to be strong entacapone accumulators, only a small percentage of *R. bromii* and part of the *B. uniformis* population accumulated entacapone (Fig. 3d,e). High entacapone signals were also detected in cells not targeted by FISH (Fig. 3e). We further confirmed the ability of *P. dorei* to accumulate entacapone when grown in pure culture (Extended Data Fig. 5). Interestingly, while entacapone accumulation drastically inhibited the growth of *P. dorei* as a microbiome community member and in pure culture, it did not affect *Streptococcus* growth in the community to the same extent and showed even growth promotion for *E. coli* (Fig. 3e). Thus, bioaccumulation of the drug led to growth inhibition for certain taxa, whereas others were unaffected or even stimulated in growth. This is in line with findings reported for the well-studied antidepressant duloxetine, which was also found to be bioaccumulated by gut microbiota taxa without impacting their growth^13^.

### Entacapone chelates iron and induces iron starvation in whole microbiome populations

Entacapone’s nitrocatechol group can act as a bidentate ligand, chelating and forming stable complexes with transition metal ions such as iron (Fe) through the catecholate oxygen atoms^40^. Using an assay widely applied to detect siderophores and other strong Fe-chelating agents in solution, we confirmed entacapone’s ability to chelate ferric iron (Fe(III)) (Fig. 4a). Binding of entacapone to metal ions such as Fe(III) has been proposed to occur via formation of a tris complex^40^. The stability constant of entacapone’s association with Fe (pFe(III)=19.3) has been demonstrated to be similar to constants described for other known iron chelators such as 2,2′-bipyridyl (pFe(III)=21.5)^41^, but lower than reported for the siderophore enterobactin (pFe(III)=49)^42^. Entacapone is also predicted to complex ferrous iron (Fe(II)) with rather high affinity, but it is not predicted to form strong complexes with any other metal cations^41^.

**Figure 4.**
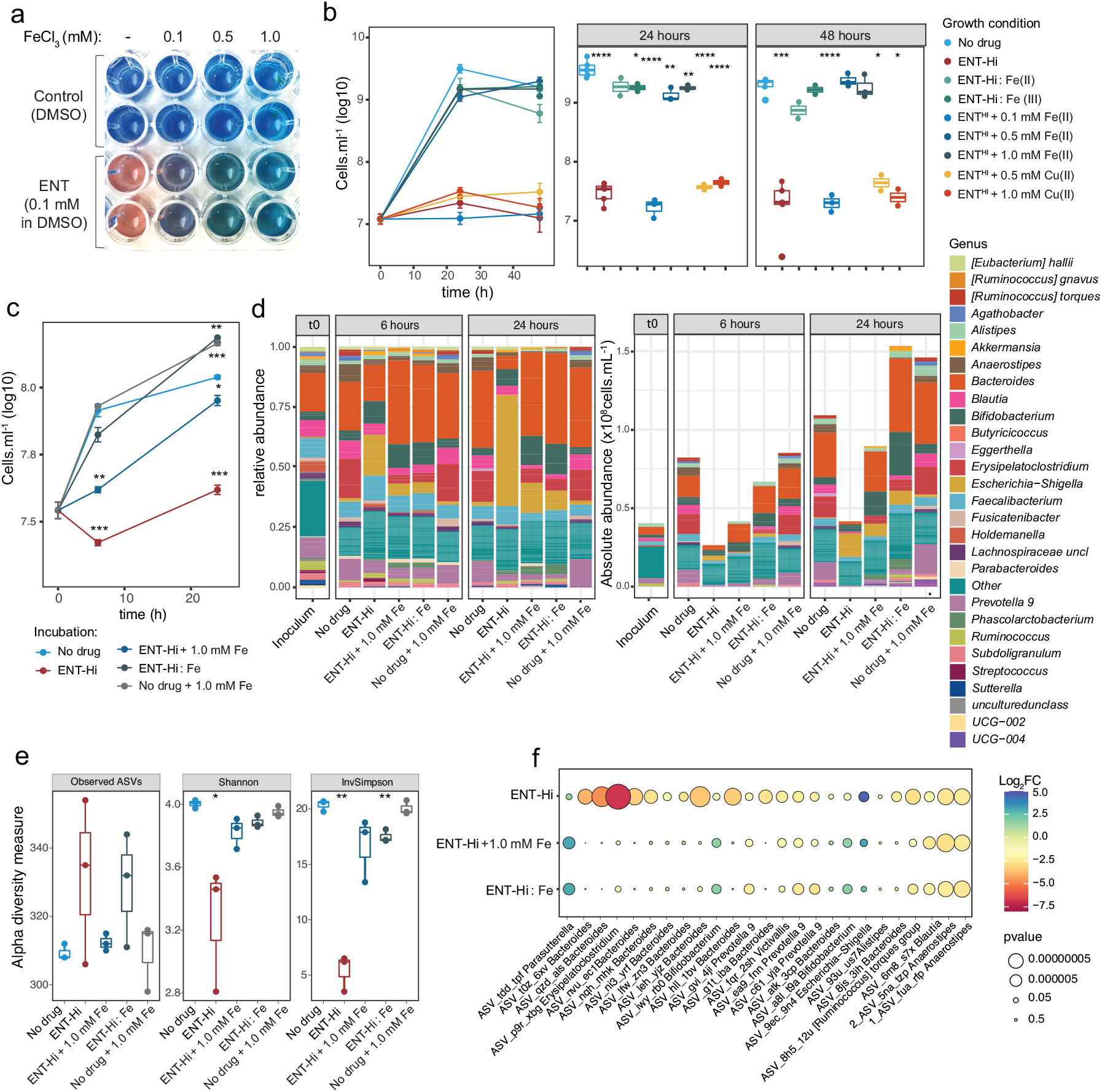
Iron supplementation rescues the impact of entacapone on the gut microbiome. **a.** Siderophore detection assay showing the change of color of the indicator complex from blue to orange/pink (see Methods) in the presence of 0.1 mM of entacapone (in DMSO), but not in the presence of DMSO alone. The indicator complex changed back to its original color (blue) after addition of an excess of ferric iron (FeCl_3_, added at 0.1 to 1.0 mM). **b.** Growth of *Bacteroides thetaiotaomicron* in the absence of drug or in the presence of: ENT-Hi; ENT-Hi preloaded with 1 mM FeSO_4_ (ENT-Hi:Fe(II)), ENT-Hi, ENT-Hi preloaded with 1 mM FeCl_3_ (ENT-Hi:Fe(III)); or ENT-Hi supplemented with the indicated concentrations of Fe(II) (FeSO_4_) or Cu(II) (CuCl_2_)(see Methods). The box plots on the right refer to the same data as displayed on the left plot, but split by the time of incubation (24 and 48 hours), for better visualization. **c.** Total cell loads in faecal samples incubated for 6 and 24 hours with either ENT-Hi, 1mM FeSO_4_ (No drug + 1.0 mM Fe), both (ENT-Hi + 1.0 mM Fe), or ENT-Hi pre-incubated with 1 mM of FeSO_4_ (ENT-Hi: Fe). Total cell loads were assessed by flow cytometry. **d.** Relative (left panels) and absolute (right panels) genus abundance profiles of faecal sample microbial communities incubated as described in c and assessed by 16S rRNA gene amplicon sequencing. All genera present at relative abundances below 0.01 or absolute abundances below 1.5×10^6^ cells.ml^-1^ were assigned to the category “Other”. **e.** Alpha diversity metrics (Observed ASVs, Shannon index and Inverse Simpson’s diversity index) in gut microbial communities described in d. **f.** Bubble plot denoting the fold change (FC, represented as Log_2_FC) in absolute abundance relative to control incubations for the 25 most abundant ASVs that are affected by ENT-Hi treatment, as assessed by DeSeq2 analyses. All ASVs have a base mean >50000, fold change of >1 or <-1 (relative no drug control), and p_adjusted_<0.05. In **b**, **e** boxes represent the median, first and third quartile. Whiskers extend to the highest and lowest values that are within one and a half times the interquartile range. In **b**, **c** and **e:** *p<0.05; **p<0.01; ***p<0.001, ****p<0.0001; unpaired two-samples t-test with “No drug” used as a reference group. Only statistically significant differences are indicated.

Iron is a limiting nutrient in the gut and essential for growth of most gut microbes^43^. Iron concentrations in stool are estimated to be around 60 μM^44^, and the medium used here for faecal sample incubations contained similar concentrations of iron (31 μM). As the estimated concentration for entacapone in the large intestine is approximately 2 orders of magnitude higher (1965 μM), we postulated that the inhibitory effect of ENT-Hi on microbial growth could be directly related to its ability to deprive gut microbes from iron via chelation, similarly to what has been documented for other Fe(II) and Fe(III) chelators. To test this hypothesis, we grew *Bacteroides thetaiotaomicron,* a gut commensal severely impacted by Ent-Hi (log2-fold change: −4.16, adjusted p-value: 0.016, Supplementary Table 5: ASV_g1t_iba) in the presence of ENT-Hi or Fe-loaded ENT-Hi (Fig. 4b). Fe-loaded entacapone was obtained by addition of 1 mM of Fe(II) or Fe(III) to ENT-Hi followed by removal of any excess iron cations via phosphate precipitation^45^ (Fig.4b; see Methods). While ENT-Hi inhibited the growth of *B. thetaiotaomicron*, ENT-Hi pre-complexed with either Fe(III) or Fe(II) did not, enabling *B. thetaiotaomicron* to grow normally (Fig.4b). This is in agreement with previous studies reporting that supplementation of the iron chelator 2,2′-bipyridyl inhibits growth of *Bacteroides fragilis,* but this effect is largely diminished when bipyridyl is saturated with Fe(II)^46^. This is because iron-saturated chelators are no longer able to complex free Fe(II) present in the medium (or intracellularly), thus enabling bacteria to access iron and grow normally. As we expected iron to be mostly present in the lower oxidation state under anaerobic conditions, we used Fe(II) in all subsequent incubations. Supplementation of ENT-Hi alone followed by addition of increasing amounts of Fe(II) also alleviated the inhibitory effect of ENT-Hi on *B. thetaiotaomicron*, but only when Fe(II) was added at concentrations of 0.5 mM or above (Fig.4b). Addition of similar amounts of cupric (Cu(II)) ions did not reverse growth inhibition by ENT-Hi (Fig.4b), confirming that the observed effect is specific for iron. In summary, these results demonstrate that supplementation of Fe-salts alone or of an ENT-Hi:Fe complex rescues the inhibitory effect of ENT-Hi on *B. thetaiotaomicron,* strongly indicating that entacapone drives iron limitation.

To determine whether the results above also apply to a complex microbiota, we incubated faecal samples under equivalent conditions as described in Fig. 1a with ENT-Hi, ENT-Hi preloaded with Fe(II) or ENT-Hi followed by addition of Fe(II). In agreement with the results obtained for *B. thetaiotaomicron* alone, supplementation of the whole microbiome with iron-loaded ENT-Hi or with ENT-Hi followed by 1 mM of ferrous iron resulted in a complete or near complete reversal of the inhibitory effect on microbial biomass accumulation (Fig. 4c,d, Supplementary Table 12). Importantly, we confirmed that this effect is not due to lower cellular uptake of iron-complexed entacapone, as ENT-Hi:Fe(II) bioaccumulates in microbiota cells to the same level or higher than ENT-Hi alone (Extended Data Fig. 6), thus suggesting that the

ENT-Hi:Fe(II) complex behaves similarly to entacapone alone in terms of its ability to penetrate and accumulate in cells. Supplementation of ferrous iron rescues the impact of ENT-Hi on the community alpha diversity metrics as well as on the growth of nearly all of the top 25 most affected taxa (exceptions are Firmicutes such as *Anaerostipes* and *Blautia*) (Fig.4e,f; Supplementary Tables 13 and 14). Iron supplementation does not only enable taxa negatively impacted by entacapone to grow, but it also seems to restrict the accelerated expansion of organisms such as *E. coli*, prompted by the presence of ENT-Hi alone (Fig.1e and Fig.4d). All together these results strongly suggest that complexation of the limiting nutrient iron by entacapone is the primary mechanism behind the strong inhibitory effect of entacapone on the microbiome.

### Entacapone promotes growth of iron-scavenging *E. coli*

Next, we interrogated the capability of the few indigenous microbiome members to thrive under the iron-limiting conditions induced by ENT-Hi. Several commensal and pathogenic *Enterobacteriaceae*, including *E. coli*, are known to synthesize and release siderophores which bind to Fe(III) with high affinity, enabling them to scavenge iron when it becomes limiting and in this way, enhance their gut colonization^47^. The most dominant organism in ENT-Hi incubations is an *E. coli* strain classified as *E. coli*_D (according to GTDB^48^), for which we were able to recover a complete MAG (bin.187, Supplementary Table 7). A search for genes involved in siderophore synthesis within the *E. coli*_D MAG led us to identify an entire gene cluster coding proteins necessary for synthesis, export, and import of the siderophore enterobactin (Fig. 5a). Thus, we postulate that the ability of *E. coli*_D to produce enterobactin enables it to grow under iron-limiting conditions induced by ENT-Hi. Indeed, after isolating this *E. coli* strain from glycerol-preserved ENT-Hi incubations (isolate E2 described in methods), we could confirm its ability to grow in minimal medium prepared without iron (Fig. 5b). We further demonstrated that its growth is not impacted (neither inhibited nor promoted) by the presence of ENT-Hi when grown under isolation (Fig. 5b). Enterobactin chelates ferric iron with much higher efficiency than entacapone^41^, and our results suggest that it enables *E. coli* to scavenge and acquire enough iron to sustain its growth under the iron-limiting conditions induced by the presence of entacapone (Fig. 5c). The Ton-B siderophore receptor FepA may also enable *E. coli* to uptake and scavenge iron from exogenous catecholate:iron complexes^49^, possibly even directly from iron-complexed entacapone. We could not find any genes involved in the production of known siderophores in the MAG of the ENT-Hi thriving *Ruminococcus*, and thus we speculate it expands in the entacapone supplemented medium by importing iron-loaded siderophores produced by *E. coli*, or by producing a yet unidentified siderophore or other high-affinity iron binding proteins (Fig. 5c).

**Figure 5.**
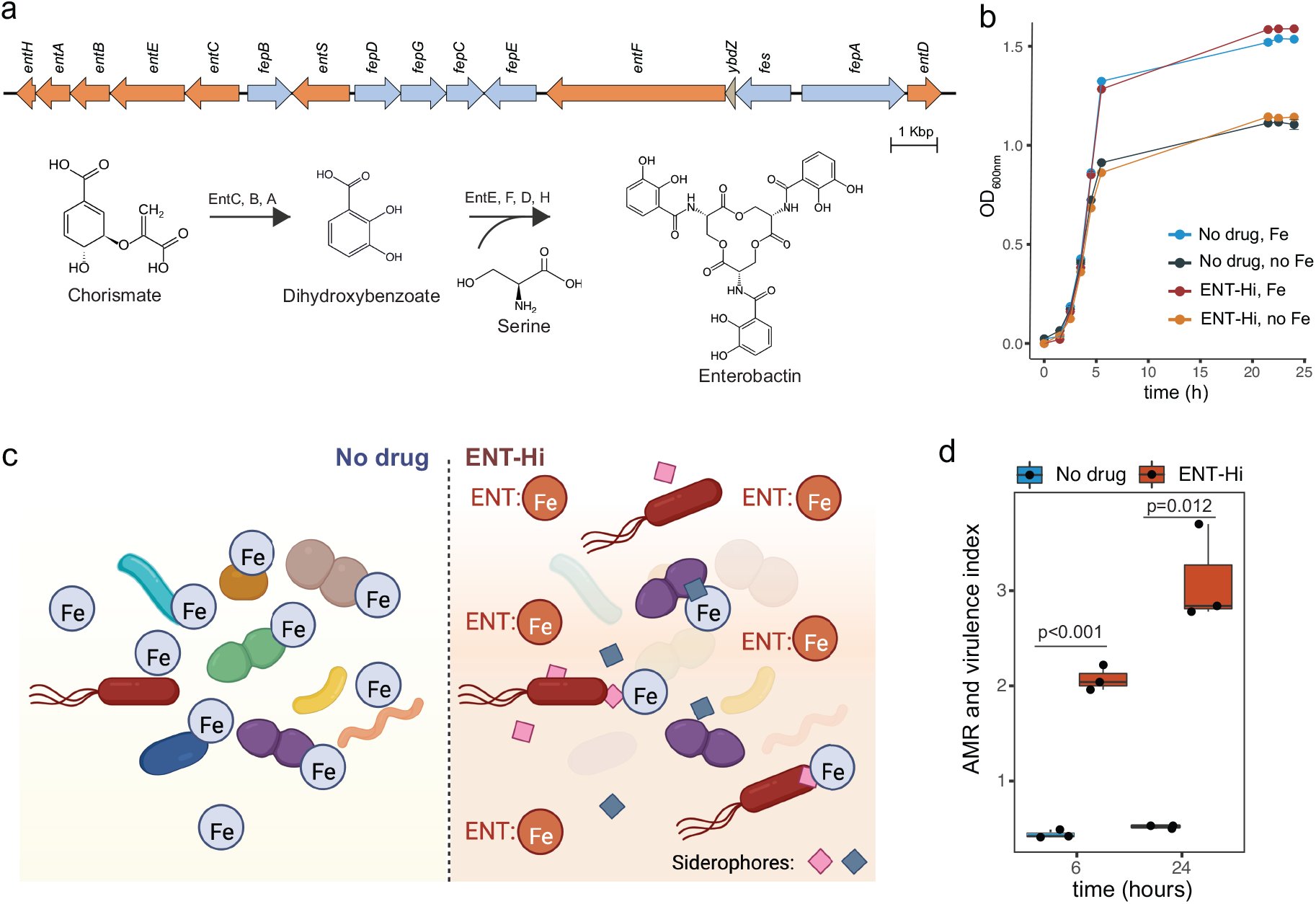
The iron-limiting conditions induced by entacapone select for siderophore producers and drive an increase in AMR and virulence potential in the faecal microbiome. **a.** Genetic organization of the enterobactin biosynthesis locus in *E. coli* D bin.187. Genes encoding enzymes involved in enterobactin biosynthesis (*entABCDEFH*) and export (*entS*) are highlighted in orange. The steps of enterobactin synthesis catalysed by the products of these genes are shown on the bottom. Genes involved in enterobactin import and iron release are highlighted in blue. **b.** Growth of an *E. coli* D bin.187 isolate determined by optical density measurements of cultures supplemented or not with 1965 μM entacapone (ENT-Hi) and/or 100 μM of FeSO_4_. **c.** Schematic representation of the working hypothesis. In the absence of entacapone (left panel), enough iron is available to most gut microbiome members. In ENT-Hi conditions, entacapone complexes available iron (ENT:Fe), and only organisms able to produce siderophores (represented with blue and pink diamonds) for iron scavenging are able to grow and thrive (right panel). **d.** Increase in the AMR and virulence index (see Methods) in no drug controls and entacapone (ENT-Hi) treated samples. Indicated p-values were obtained from unpaired two-sample t-tests.

Siderophore production and iron scavenging are commonly observed in pathogenic or pathobiont strains that also tend to encode other virulence traits^50^. A screening of our MAG catalog for the presence of virulence factors and antimicrobial resistance genes^51^ identified a diversified panel of AMR and virulence genes to be present in the retrieved gut MAGs (Supplementary Table 15), even though these samples originated from healthy individuals. These include genes encoding resistance to, among others, beta-lactams, tetracyclines, macrolides and aminoglycosides, as well as more general antimicrobial resistance genes such as efflux pumps, and virulence genes. ENT-Hi drives an increase in the abundance of AMR and virulence within faecal microbiomes (Fig. 5d). This increase is driven in great part by the increase in abundance of *E. coli_D*, whose genome encodes a total of 14 AMR and virulence genes, in addition to at least one siderophore production cluster, thus suggesting a pathogenic potential of this organism (Supplementary Table 15). Overall, these results reveal that ENT-Hi promotes the growth of iron scavenging organisms with an associated pathogenic potential.

## Discussion

Gaining a deeper insight into the interactions between drugs and the gut microbiome is essential for revealing and predicting how the microbiome might influence the availability, efficacy and toxicity of pharmaceuticals. Here, we evaluated the impact of the nervous-system targeted drugs loxapine succinate and entacapone on whole microbiomes derived from faecal samples. Both drugs caused major shifts in microbial communities at physiologically relevant concentrations, inhibiting the growth of many taxa while stimulating others. Using single-cell activity measurements, we further revealed fine changes in the activity of specific community members even at low drug concentrations and that were not captured by commonly used 16S rRNA gene sequencing and abundance profiling. For instance, we did detect reduced *B. uniformis* and *Clostridium* sp900539375 activities at 6 hours, but their abundance is not affected until 24 hours (Fig. 1e, Fig 2d). This is a major advantage of our approach as it captures drug-induced changes in short incubation times, which are ideal for *ex vivo* systems like the one in use. Importantly, our results show that loxapine succinate exerts an effect on a broader range of taxa in the context of the community than its effects on microbes grown under isolation^1^. This highlights the necessity of using complex microbial communities for a better assessment of drug-microbiota interactions. One way by which some taxa may sensitize or protect others to a particular drug is through chemical conversion or accumulation of the drug^10^. However, previous studies have shown that the gut microbiota does neither significantly bioaccumulate nor transform loxapine succinate^10^. Thus, we presume that cross-sensitization to loxapine is likely due to drug-induced changes in microbial metabolites that are involved in interspecies interactions.

The gut microbiome produces metabolites that can signal to the host and influence several aspects of host physiology, including brain function^52^. Recent studies suggest that modulation of the microbiome by drugs may contribute to the therapeutic effect of antipsychotics^1^. Interestingly, our results demonstrate that loxapine succinate promotes the growth of several key gut species, including *Lachnospiraceae* (e.g. *Anaerostipes*) and *Butyricicoccaceae* species, which have been shown to produce neuroactive metabolites such as short chain fatty acids (SCFAs)^53^. SCFAs might directly or indirectly communicate along the microbiota-gut-brain axis by activating G protein-coupled receptors or inhibiting histone deacetylases^54^. Additionally, they can modulate the blood-brain barrier, activate the vagus nerve, facilitate the secretion of other hormones or neurotransmitters, as well as interfere with the immune response^54^, all these factors contributing to its neuromodulatory effects. Thus, it may be plausible to consider changes in the microbiome as an additional mode of action of loxapine succinate. This may also help to explain why it may take weeks for this antipsychotic to act entirely^55^, as its therapeutic effects may first require changes in the gut microbiome.

The impact of entacapone on the microbial community was very pronounced, selecting for the growth of organisms with pathogenic potential, such as *E. coli*. An increased abundance of *Enterobacteriaceae* was previously reported for Parkinson’s disease (PD) patients taking entacapone^56^, while other studies reported increases in *Enterococcaceae, Bifidobacteriaceae*, *and Clostridiales* family X^57^. These inconsistencies across studies can be partially attributed to the cross-sectional and rather small number of patients involved in each study. An increase in *Enterobacteriaceae* has also been linked to the side effects of other drugs^58^ and may also help to explain diarrhoea experienced by some patients taking entacapone. In addition, entacapone is always prescribed in combination with levodopa to inhibit its off-site metabolism, and frequently also in combination with carbidopa, which inhibits levodopa off-site decarboxylation^27^. While levodopa has been shown to exert no effect on the distal gut microbiota, probably due to its high absorption in the upper gastrointestinal (GI) tract, future studies would be valuable to investigate the effects of a simultaneous supplementation of entacapone and levodopa to whole microbiome communities, especially in the upper GI tract. Biotic conversion of levodopa to dopamine by *Enterococcus* spp. and *Lactobacillus* spp. tyrosine decarboxylases in the small intestine significantly reduces its bioavailability to the host^59^. Given the strong inhibitory effect of entacapone on the microbiota, it would be interesting to evaluate whether entacapone can inhibit taxa that metabolize levodopa in the upper gastrointestinal tract, thereby contributing also in this way to increase the bioavailability of levodopa.

Like levodopa, entacapone has also been shown to be metabolized by gut microbial species^10^. Here, we demonstrate that entacapone is also bioaccumulated, with *Bacteroides, Phocaeicola, Streptococcus* and *Escherichia* spp. being able to accumulate sufficient amounts of entacapone for photothermal detection (Fig. 3). This accumulation likely results in depletion of entacapone in the surrounding environment overtime, thus explaining the slight alleviation of entacapone’s inhibitory effect between 6 and 24 hours of incubation, when compared to the first 6 hours (Fig. 1b). Interestingly, using single-cell chemical imaging we further show that this bioaccumulation is heterogeneous, with some cells within a particular taxon accumulating varying amounts of the drug. In addition, we further show that entacapone bioaccumulation does not necessarily lead to growth inhibition (Fig. 3e, f), similarly to what has been described before for other bioaccumulated drugs. It remains to be determined whether entacapone bioaccumulation can be linked to its metabolism. Bacterial metabolism of entacapone occurs mostly by means of nitroreduction, which is prone to occur under anaerobic conditions, with several *Bacteroides* spp. encoding genes linked to entacapone nitroreduction^10^. Given the extensive conversion of entacapone by gut taxa observed previously, we predict a large fraction of entacapone’s nitrocatechol group to be in its reduced form in our incubations and in the gut. Despite its bioaccumulation and/or conversion, we demonstrate that entacapone exerts a strong effect on the gut community due to its ability to complex iron via the catechol group.

Iron represents an essential enzyme cofactor in most bacteria^43^. Iron complexation by entacapone led to a decrease in biomass accumulation and in the abundance of most of the top abundant taxa detected in these samples, an effect that was reversed by supplementation of iron (Fig. 4c-f). Taxa not found to be significantly impacted by entacapone were presumably able to grow by relying on intracellularly accumulated iron or on high-affinity surface-associated iron transporters that are ubiquitous in bacteria. Among stimulated taxa, the siderophore-producing *E. coli*_D strain present in our incubations greatly benefitted from entacapone’s presence, but only in the context of the community, as entacapone supplementation to the isolate alone did not cause any significant boost in *E.coli*_D growth (Fig. 5b). Thus, taxa stimulated by entacapone likely acquired iron via the mechanisms mentioned above, or via siderophores, and expanded in the community at the cost of nutrients released by dead cells or supplied via cross-feeding from other species. Treatment of communities with other iron complexing agents has shown somewhat similar effects on the gut community^60,61^.

The expansion of an organism with siderophore synthesis, AMR and virulence potential in the presence of entacapone is concerning. Most successful gut pathogens tend to encode siderophore production systems and are therefore also expected to take advantage of the entacapone-induced remodeling of the gut community. It would be important to determine if entacapone treatment increases the likelihood of intestinal infections, similarly to what has been observed in patients taking PPIs^19^. As oral iron supplementation reduces levodopa and entacapone absorption in the small intestine^62,63^, the effect of entacapone on the microbiome could be circumvented in the future by delivering iron specifically to the colon of patients taking the drug. If administered in appropriate amounts, this new adjuvant therapy could be expected to aid in preservation of microbiome homeostasis for patients taking entacapone, and presumably also for patients taking other catechol-containing drugs that might reach relevant concentrations in the large intestine (e.g. opicapone, apomorphine, fenoldopam mesylate) and that might affect the microbiome via similar mechanisms. Colon-targeted adsorbents, such as DAV132, could also be tested, as these have shown the ability to sequester antibiotic drugs and reduce antibiotic-induced effects on the microbiome^64^. Our results advance our understanding of the impact of antipsychotic and antidyskinetic drug therapies on microbiome homeostasis, their mechanisms of action, and can direct future optimization of such therapies.

## Methods

### *Ex-vivo* gut microbiome incubations with drugs

Human faecal samples were collected from six healthy adult individuals (two males and four females between the ages of 25 to 39) who had not received antibiotics in the prior 3 months. Study participants provided informed consent and self-sampled using an adhesive paper-based feces catcher (FecesCatcher, Tag Hemi, Zeijen, NL) and a sterile polypropylene tube with the attached sampling spoon (Sarstedt, Nümbrecht, DE). The study protocol was approved by the University of Vienna Ethics Committee (reference #00161). All meta(data) is 100% anonymized and compliant with the University’s regulations. Samples were transferred into an anaerobic tent (Coy Laboratory Products, USA) within 30 min after sampling, and all sample manipulation and incubations were performed under anaerobic conditions (5% H2, 10% CO2, 85% N2). Each sample was suspended in M9 mineral medium supplemented with 0.5 mg.mL^-1^ D-glucose (Merck), 0.5% v/v of vitamin solution (DSMZ Medium 461) and trace minerals, herein referred to as sM9. Samples were suspended in sM9 to yield a 0.05 g.mL^-1^ faecal slurry. At this point one aliquot of each sample was collected, pelleted, and stored at −80°C for metagenomic analysis. The homogenate was left to settle for 10 minutes, and the supernatant (devoid of any large faecal particles) was transferred into a new flask, where supernatants from the six different donors were combined. This combined sample was further diluted 1:10 in sM9 medium (as described above) or in supplemented Brain Heart Infusion (BHI) medium containing either 0% or 55% D2O (99.9% atom % (at%) D; Merck) for final 0% (control) or 50% D2O in incubation medium (Fig. 1a). Supplemented BHI medium consisted of 37 g.L^-1^ of brain heart infusion broth (Oxoid), 5 g.L^-1^ yeast extract (Oxoid), 1 g.L^-1^ L-cysteine (Merck) and 1 g.L^-1^ NaHCO3 (Carl Roth GmbH, Germany). Incubation tubes were supplemented with dimethylsulfoxide (DMSO, from Merck), entacapone (Prestwick Chemicals) or loxapine succinate (Prestwick Chemicals) pre-dissolved in DMSO. The final concentration was 2% w/v of DMSO in all vials (except for the H2O control, where water was added instead of DMSO). A subset of vials was supplemented with 20 μM or 1965 μM entacapone (ENT-Low and ENT-Hi, respectively) and another subset was supplemented with 20 μM or 100 μM loxapine succinate (LOX-Low and LOX-Hi, respectively). The colon concentration estimated for entacapone is 1965 μM^1^, while for loxapine no estimate was available. We predicted that loxapine would reach similar colon concentrations as its chemical and therapeutic analogues amoxapine and clozapine, which are estimated to reach colon concentrations of 138 μM and 153 μM, respectively^1^. Using these values as a reference, we predict that loxapine succinate should be present in the colon at concentrations of at least 100 μM and chose it as the LOX-Hi concentration. At time 0, and after an incubation time of 6 or 24 hours at 37°C under anaerobic conditions, two sample aliquots from each incubation and controls were collected by centrifugation. One aliquot was washed with 1× PBS and then fixed in 3% paraformaldehyde solution for 2 h at 4°C. Samples were finally washed two times with 1 ml of PBS and stored in PBS:Ethanol (50% v/v) at −20°C until further use. The second pelleted aliquot was stored at −20°C until further processing. A third aliquot was collected into sealed anaerobic vials containing 40% glycerol (Carl Roth GmbH, Germany) in PBS for a final cell 50% v/v cell suspension in 20% glycerol and stored at −80°C until further use.

For iron rescue experiments, fresh faecal samples received from the same individuals (except for one male that was traveling at the time of the experiment) were collected and processed as described above. To establish appropriate controls for imaging of entacapone bioaccumulation by microbiota cells, an aliquot of the freshly prepared 0.05 g.mL^-1^ faecal slurry was immediately fixed with either 3% PFA solution or ethanol (50% v/v) at 4°C for 2 hours. Fixed faecal samples were washed with 1xPBS as described above and incubations with fixed samples and entacapone or entacapone:iron (see below) were conducted in parallel with incubations using live samples. Incubation vials were then supplemented with DMSO 2% v/v with or without 1965 μM entacapone, in the presence or absence of supplemented iron (1mM FeSO4, Merck; Fig. 5). Additional incubation vials (triplicates) were treated with entacapone pre-complexed with iron: briefly, entacapone and iron (FeSO4 or FeCl3) powder were mixed and resuspended in 2 mL of DMSO yielding a final concentration of 1965 μM entacapone and 1 mM FeSO4 (or FeCl3) and stored overnight under anaerobic conditions. The next day, 120μL of sodium phosphate monobasic (0.5M) was added and the samples were mixed well. After 20 minutes, samples were centrifuged for 5 min, 14000 *g* to remove unbound iron precipitated by the addition of sodium phosphate^45^. The supernatant containing the iron-complexed entacapone was collected into a new eppendorf tube and supplemented to the faecal incubation vials. Incubations were sampled as described above.

### Cell counts from *ex-vivo* microbiome incubations

Microbial loads in faecal incubation vials were determined using flow cytometry and counting beads as detailed below. Samples preserved in glycerol were diluted 200 to 800 times in 1x PBS (Supplementary Table 1). To remove any additional debris from the faecal incubations, samples were transferred into a flow cytometry tube by passing the sample through a snap cap containing a 35 μm pore size nylon mesh. Next, 500 μL of the microbial cell suspension was stained with the nucleic acid dye SYTO™ 9 (Thermo Fisher Scientific, 0.5 μM in DMSO) for 15 min in the dark. The flow cytometry analysis of the microbial cells present in the suspension was performed using a BD FACSMelody™ (BD Biosciences), equipped with a BD FACSChorus™ software (BD). Briefly, background signals from the instrument and the buffer solution (PBS) were identified using the operational parameters forward scatter (FSC) and side scatter (SSC). Microbial cells were then displayed using the same settings in a scatter plot using the forward scatter (FSC) and side scatter (SSC) and pre-gated based on the presence of SYTO™ 9 signals. Singlets discrimination was performed. Absolute counting beads (CountBright^TM^, ThermoFisher Scientific) added to each sample were used to determine the number of cells per mL of culture by following the manufacturer’s instructions. Fluorescence signals were detected using the blue (488 nm – staining with SYTO™ 9 and CountBright™ beads) and yellow-green (561 nm - CountBright^TM^ beads only) optical lasers. The gated fluorescence signal events were evaluated on the forward–sideways density plot, to exclude remaining background events and to obtain an accurate microbial cell count (Supplementary Table 1). Instrument and gating settings were identical for all samples (fixed staining–gating strategy).

### Nucleic acid isolation and 16S rRNA gene amplification and sequencing

Pellets of microbiome incubation samples were resuspended in 600 μl of lysis solution RL (InnuPREP DNA/RNA mini kit, Analytik Jena) and subjected to bead beating for 30 seconds at 6.5 m/s in lysis matrix E (MPBiomedicals) tubes. After pelleting cell debris for 10 minutes at 8000g, supernatants were transferred into the InnuPREP DNA/RNA mini kit spin filter tubes (Analytik Jena) and DNA and RNA were extracted according to the manufacturer’s protocol. Amplification of bacterial and archaeal 16S rRNA genes from DNA extracts was performed with a two-step barcoding approach^65^ (UDB-H12).

In the first-step PCR, the primers 515F66 (5′-GTGYCAGCMGCCGCGGTAA-3′) and 806R67 (5′-GGACTACNVGGGTWTCTAAT-3′), including a 5′-head sequence for 2-step PCR barcoding, were used. PCRs, barcoding, library preparation and Illumina MiSeq sequencing were performed by the Joint Microbiome Facility (Vienna, Austria) under project numbers JMF-2208-05 and JMF-2103-29. First-step PCRs were performed in triplicate (12.5 μl vol per reaction) with the following conditions: 1X DreamTaq Buffer (Thermo Fisher), 2 mM MgCl2 (Thermo Fisher), 0.2 mM dNTP mix (Thermo Fisher), 0.2 μM of forward and reverse primer each, 0.08 mg ml−1 Bovine Serum Albumin (Thermo Fisher), 0.02 U Dream Taq Polymerase (Thermo Fisher), and 0.5 μl of DNA template. Conditions for thermal cycling were: 95°C for 3 min, followed by 30 cycles of 30 s at 95°C, 30 s at 52°C and 50 s at 72°C, and finally 10 min at 72°C. Triplicates were combined for barcoding (with eight PCR cycles). Barcoded samples were purified and normalized over a SequalPrep Normalization Plate Kit (Invitrogen) using a Biomek NXP Span-8 pipetting robot (Beckman Coulter), and pooled and concentrated on PCR purification columns (Analytik Jena). Indexed sequencing libraries were prepared with the Illumina TruSeq Nano Kit as described previously68, and sequenced in paired-end mode (2× 300 bp; v3 chemistry) on an Illumina MiSeq following the manufacturer’s instructions. The workflow systematically included four negative controls (PCR blanks, i.e., PCR-grade water as template) for each 90 samples sequenced. The 16S rRNA gene sequences were deposited in the NCBI Sequence Read Archive (SRA) as BioProject Accession PRJNA1033532.

### Analysis of 16S rRNA gene amplicon sequences

Amplicon pools were extracted from the raw sequencing data using the FASTQ workflow in BaseSpace (Illumina) with default parameters^65^. Demultiplexing was performed with the python package demultiplex (Laros JFJ, github.com/jfjlaros/demultiplex) allowing one mismatch for barcodes and two mismatches for linkers and primers. DADA2^69^ R package version 1.16.0 (https://www.r-project.org/, R 4.0.2) was used for demultiplexing amplicon sequencing variants (ASVs) using a previously described standard protocol^70^. FASTQ reads were trimmed at 150 nt with allowed expected errors of 2. Taxonomy was assigned to 16S rRNA gene sequences based on SILVA taxonomy^71^ (release 138) using the DADA2 classifier.

Samples were analyzed using the vegan (v2.5-.6; https://CRAN.R-project.org/package=vegan) and phyloseq^72^ (v1.30.0) packages of the software R (https://www.r-project.org/, R 4.0.2). For samples subjected to different drug treatments, sequencing in parallel of two extraction controls (without adding faecal samples) yielded 10 (control 1) and 189 reads (control 2). In control 1, 9 of the 10 reads were assigned to Cyanobacteria or chloroplast and were not detected in the samples. Likewise, in control 2, more than 90% of the reads originated from either Cyanobacteria ASVs or a single Comamonadaceae (*Aquabacterium*) not detected in any of the samples. These ASVs were removed from analysis. The remaining negative control reads (control 1: 1 read, control 2: 8 reads) were assigned to taxa typically found in the gut that were also detected in the samples and were therefore retained for subsequent analyses. We assume these low number of reads to originate from a low level of cross-contamination that can occur when multiple samples are handled in parallel. After quality filtering and removal of contaminant sequences, a total of 1132 ASVs were retained. The average read number per sample was 11176 ± 3087 high-quality sequences, and sample coverage was above 98% (Supplementary Table 2). For alpha and beta diversity analysis, sequence libraries were rarefied to 4681 reads per sample. For samples referring to the entacapone and iron supplementation experiment, sequencing in parallel of two extraction controls yielded 2 (control 1) and 134 reads (control 2). After quality filtering and removal of contaminant sequences (using the rationale described above), a total of 716 ASVs were retained. The average read number per sample was 18733 ± 4427 high-quality sequences and the sample coverage was above 99% (Supplementary Table 2). For alpha diversity analysis, sequence libraries were rarefied to 9729 reads per sample. For quantitative microbiome analyses, relative abundances of each taxon in a sample were calculated after correcting for the different number of copies of the 16S rRNA gene, according to *rrn*DB (version 5.7). For this correction, we classified ASVs using DADA2 and the RDP^73^ taxonomy 18, release 11.5 (https://doi.org/10.5281/zenodo.4310151), by applying default parameters. These corrected relative abundances were then multiplied by the total microbial loads obtained from flow cytometry (Supplementary Table 1), yielding the total abundance of each taxon per sample.

DESeq2^74^ (v1.26.0) implemented in phyloseq was used to identify significant differences in ASV abundances between drug treatments. Only ASVs that had in total ≥10 reads (relative abundance microbial profile) or 5.0e5 reads (quantitative microbial profile) were considered for comparisons by DESeq2 analyses. All statistical analysis of microbiome data was carried out with the software R (R 4.0.2). The applied significance tests and obtained p-values are referred to in the main text and figure legends.

### Long-read sequencing

DNA for long-read sequencing was isolated using the DNeasy PowerSoil Pro Kit (Qiagen), according to the manufacturer’s instructions. A pool of 6 DNA extracts was prepared for sequencing using the ligation sequencing kit (SQK-LSK112, Oxford Nanopore Technologies) following the manufacturer’s protocol. The DNA was sequenced on a Promethion P24 (Oxford Nanopore Technologies) on a R10.4 flowcell (FLO-PRO112, Oxford Nanopore Technologies). The DNA sequencing was carried out using Minknow (v. 21.10.8, Oxford Nanopore Technologies).

### Shotgun metagenomic sequencing

The same 6 samples were individually sequenced in an Illumina Novaseq 6000 platform by the Joint Microbiome Facility (Vienna, Austria), under project number JMF-2110-04. The illumina reads were trimmed using cutadapt^75^ (v. 3.1). Illumina reads were mapped to the assemblies using Minimap2^76^ (v. 2.17).

### Metagenomic analysis

The Nanopore reads were assembled using flye^77^ (v. 2.9-b1768) with “–nano-hq,” polished three times with Minimap2^76^ (v. 2.17) and Racon^78^ (v. 1.4.3), followed by two rounds of polishing with Medaka (v. 1.4.4, github.com/nanoporetech/medaka). Illumina and Nanopore reads (Supplementary Table 6) were mapped to the assemblies using Minimap2 (v. 2.17) and read mappings were converted using SAMtools^79^ (v. 1.12). Read coverage and automatic binning was performed using MetaBAT2^80^ (v. 2.15). Contigs labeled as circular by the assembler were extracted as independent bins before the automated binning process. The quality of the recovered metagenome-assembled genomes (MAGs) was checked using QUAST^81^ (v. 5.0.2) and CheckM^82^ (v. 1.1.1), and genomes were classified using GTDBtk^48^ (v. 1.5.1). rRNA genes were detected using Barrnap (v. 0.9, https://github.com/tseemann/barrnap.) and tRNA genes were detected using trnascan^83^ (v. 2.0.6). MAGs with a completeness >90 % but where barrnap did not pick up a 5S rRNA gene were checked for 5S rRNA genes using INFERNAL^84^ (v. 1.1.3).

All MAGs were searched for anti-microbial resistance (AMR) and virulence genes using AMRFinderPlus^51^ (v.3.10.21). The AMR and virulence index (Fig. 5a) was calculated as follows: the total copies of AMR and virulence genes found to be present in each MAG were multiplied by the absolute abundance of the MAG (abundance of the ASV matching the 16S rRNA gene of the MAG) in the sample. The same was repeated for all MAGs for which AMRFinderPlus identified AMR or virulence genes and by summing these we were able to predict the total number of copies of AMR and virulence genes for each sample, per mL of culture. The resulting values were then normalized to the total biomass per mL of each sample in order to obtain an AMR and virulence index per sample.

### FISH probe design and optimization

Phylogenetic analysis and FISH probe design were performed using the software ARB v. 7.0^85^. By analysis of the 16S and 23S rRNA gene, phylogenetic trees were calculated with IQ-TREE^86^ (v 1.6.12) using the RAxML GTR algorithm with 1,000 bootstraps within ARB. For abundant groups, 4 FISH probes were designed for this study and 2 additional published probes were used (Supplementary Table 9). The probes were validated *in silico* with mathFISH to test the *in-silico* hybridization efficiency of target and non-target sequences. The number of non-target sequences was assessed using the probe match function in ARB and the mismatch analysis function in mathFISH. All probes were purchased from biomers (Biomers.net, Ulm, Germany) and were double labeled with indocarbocyanine (Cy3) or sulfo-cyanine5 (Cy5) fluorochromes.

Pure cultures of *Escherichia coli* K-12, *Phocaeicola dorei* 175 (DSM17855) and *Bacteroides thetaiotaomicron* VPI-5482 (DSM2079) were grown in supplemented BHI until the mid-exponential phase and harvested by centrifugation. Pure cultures were fixed for 2 h by addition of 3 volumes of 4% (w/v) paraformaldehyde solution at 4 °C. After washing once with PBS, cells were stored in a 1:1 mixture of PBS and 96% (v/v) ethanol at − 20 °C. Where pure cultures were not available, fixed faecal samples with a high relative abundance (as determined by amplicon sequencing) of the specific target taxon was used.

To evaluate probe dissociation profiles, cells obtained from fixed pure cultures or faecal incubation samples (Supplementary Table 9) were spotted onto microscopy slides (Paul Marienfeld EN). FISH was performed as described before^87,88^, with 3 or 5h hybridization to obtain fluorescence signals with sufficient intensity. The optimal hybridization formamide concentration was found using formamide dissociation curves, obtained by application of formamide concentration series in the range from 0 to 70% in 5% increments^89^. After a stringent washing step and counterstaining using 4′,6-diamidino-2-phenylindole, samples were visualized using a Leica Thunder Epifluorescence microscope with an APO 100x/1,40 Leica oil immersion objective. Probe EUB338^90^, which is complementary to a region of the 16S rRNA conserved in many members of the domain Bacteria, was used as a positive control and a nonsense NON-EUB probe was applied to samples as a negative control for binding. Images for inferring probe dissociation profiles were recorded using the same microscopy settings and exposure times. The probe dissociation profiles were determined based on the mean fluorescence signal intensities of at least 100 probe-labeled cells and evaluated by the ImageJ software (v 1.53t). From the calculated average values, a curve was plotted and the respective value right before a decline on each curve was defined as the optimal formamide concentration.

### FISH in solution

Fixed cells (100 μl) were pelleted at 14000 *g* for 10 min, resuspended in 100 μl 96% analytical grade ethanol and incubated for 1 min at room temperature for dehydration. Subsequently, the samples were centrifuged at 14000*g* for 5 min, the ethanol was removed, and the cell pellet was air-dried. For SRS-FISH analysis, cells were hybridized in solution (100 μl) for 3 h at 46°C. The hybridization buffer consisted of 900 mM NaCl, 20 mM TRIS HCl, 1 mM EDTA, 0.01% SDS and contained 100 ng of the respective fluorescently labelled oligonucleotide as well as the required formamide concentration to obtain stringent conditions (Supplementary Table 9). After hybridization, samples were immediately transferred into a centrifuge with a rotor pre-heated at 46°C and centrifuged at 14000 *g* for 15 min at maximum allowed temperature (40°C), to minimize unspecific probe binding. Samples were washed in a buffer of appropriate stringency for 15 min at 48°C, cells were centrifuged for 15 min at 14 000 g and finally resuspended in 20 μl of PBS. Cells (5 μl) were spotted on Poly-L-lysine coated cover glasses No. 1.5H (thickness of 170 μm ± 5 μm, Paul Marienfeld EN) and allowed to dry overnight at 4°C under protection from light. Salt precipitates were removed by dipping the coverslips 2× in ice-cold Milli-Q water and allowed to dry at room temperature under protection from light.

### Picosecond stimulated Raman scattering (SRS) with widefield fluorescence microscopy

An 80-MHz pulsed laser (InSight DeepSee+; Spectra-Physics) emitting two synchronized femtosecond beams was used (Extended Data Fig. 2a). One beam was tunable in wavelength from 680 nm to 1300 nm, while the other beam had a fixed wavelength of 1040 nm. The time delay between single pulses of the two beams is adjustable by a motorized delay line on the 1040 nm beam. To implement the picosecond stimulated Raman scattering (SRS) (Extended Data Fig. 2), the fixed beam (termed Stokes beam) is intensity modulated at 2.5 MHz by an acousto-optic modulator (1205c; Isomet Corporation) and co-aligned with the tunable beam (termed pump beam), by a dichroic mirror (DMLP1000; Thorlabs). Both beams are chirped by SF57 rods to two picoseconds pulse-width and directed towards the lab-built upright microscope frame. Then, a four-focal system and a flip mirror conjugate a pair of galvo mirrors to the back aperture of a 60× water objective (UPlanApo 60XW, numerical aperture = 1.2; Olympus) or a 100× oil objective (UAPON 100XOTIRF, numerical aperture = 1.49; Olympus), allowing the collinear pump and Stokes beams to raster-scan the sample via synchronized movement of the galvo mirrors. A 1.4 numerical aperture oil condenser (Aplanat Achromat 1.4; Olympus) collects the output beams, which are then reflected by a flip mirror and filtered by a short pass filter (DMSP950; Thorlabs). Finally, the filtered-out pump beam is focused onto a silicon photodiode connected to a resonant amplifier effective at a resonant frequency around 2.5 MHz. The output alternative current signal is further amplified by a lock-in amplifier (UHFLI; Zurich Instrument) at the frequency and in phase (x channel detection) with the modulation. The output direct current signal is recorded for normalization. A data acquisition card (PCIe-6363; National Instruments) collects the output signal for image generation.

To perform widefield fluorescence imaging for FISH-visualization of the identical sample areas analyzed by SRS and photothermal imaging, two flip mirrors were flipped off (Extended Data Fig. 2a). A halogen lamp (12V100WHAL; Olympus) provides Kohler illumination of the sample from the condenser side. Then, the objective and the tube lens conjugate the sample plane to the camera (CS505CU; Thorlabs). To enable imaging of various fluorophores, different excitation and emission filter sets were inserted between lamp and condenser, and in front of the camera. For Cy3 imaging, two 530/10 nm bandpass filters (FBH530-10; Thorlabs) were used as excitation filters and two 570/20 nm bandpass filters (ET570/20x; Chroma) were used as emission filters. For Cy5 imaging, two 640 nm bandpass filters (FBH640-10; Thorlabs) were used as excitation filters and two 670/20 nm bandpass filters (ET670/50m; Chroma) were used as emission filters.

### Chemical imaging of entacapone accumulation

By utilizing the multiphoton absorption of entacapone, we detected the photothermal signal originating from optical absorption to generate entacapone distribution maps. The experimental setup was identical with picosecond SRS, but with detection of the lock-in signal by the y channel, which exhibits a π/2 phase delay relative to the intensity modulation by the AOM (Extended Data Fig. 4b). With this orthogonal phase detection, the interference of the photothermal signal with the signals emerging from cross-phase modulation and SRS was minimized (Extended Data Fig. 4d).

### Image acquisition and processing

Samples were prepared by drying fixed bacterial cells spotted onto poly-L-lysine coated coverslips (VistaVision cover glasses, No. 1; VWR) in a 4°C refrigerator and subsequent dipping into water three times to dissolve precipitates from the growth media. Then the bacteria were immersed in 5 μL of water and sandwiched by another coverslip with a 0.11 mm thick spacer in between. To acquire deuterium incorporation profiles of microbiome members labeled by FISH probes, widefield fluorescence was performed first. For different fluorophores, corresponding excitation and emission filters were applied. The signal and color gain of the camera were set to 5. Then the exposure time was adjusted to 0.5 to 5 seconds depending on the fluorescence signal intensity. The widefield transmission image was acquired by minimizing the condenser aperture and removing the filters.

To acquire the deuterium incorporation profile of the FISH-visualized cells, two flip mirrors were inserted into the beam path to guide the pump and Stokes lasers to the sample. Three SRS images, specific for Raman active vibrational modes of carbon-deuterium (C-D) bonds, carbon-hydrogen (C-H) bonds as well as the off-resonance background signal were recorded by tuning the wavelength of the pump beam to 849 nm, 796 nm and 830 nm, respectively. These wavelength values correspond to spectral wavenumbers of 2163 cm^-1^ (C-D), 2947 cm^-1^ (C-H) and 2433 cm^-1^ (silent region). Signal intensities were accumulated over increments of 20 cm^-1^. Images were acquired sequentially within the identical field of view of 32 x 32 μm^2^ with a raster step size of 106.8 nm. The per-pixel dwell time was set to 10 μs and, depending on the signal intensity level, 1∼10 image cycles were recorded to achieve a signal-to-noise ratio (SNR) of >5 for single bacterial cells in the C-H spectral region.

For acquisition of entacapone distribution maps, the pump laser was tuned to 849 nm and the signal detection was switched to the y channel of the lock-in amplifier. All images were recorded utilizing the identical scanning parameters as applied for SRS.

To process the image data sets, firstly, the illumination patterns were corrected for both widefield images (FISH) and point-scan images (SRS and entacapone distribution). Then, the widefield images and point-scan images were co-localized via a calibrated projective transform matrix. The fluorescence images were utilized to generate a single cell mask for inference of the single cell activity and the drug accumulation level. The single cell activity is expressed as %CDSRS = (ICD-Ioff)/(ICD+ICH-2Ioff), where the symbols ICD, ICH and Ioff refer to the SRS signal intensities detected at the spectral positions assigned to C-D bonds, C-H bonds and the silent region (off-resonance background). All intensity values were normalized to the direct current intensity level detected at the photodiode. The relative entacapone accumulation level is expressed as the signal intensity level detected in the y channel of the lock-in amplifier. intensity outliers (>mean±2 standard deviations) observed in the SRS signals and the photothermal signal of the human gut microbiome samples, were rejected from the single cell masks. All imaging data analysis was performed with CellProfiler and Matlab.

### Siderophore assay

The iron binding capacity of entacapone was tested using the SideroTec Assay^TM^ (Accuplex Diagnostics, Ireland), a colorimetric test for the detection of siderophores, according to the manufacturer’s instructions. Wells were read photographically and on a microplate reader at 630 nm wavelength (MultiskanTM GO Microplate Spectrophotometer, Thermo Fisher Scientific).

### Isolation and sequencing of *Escherichia coli* human gut isolate

*Escherichia coli* isolate E2 was isolated from glycerol-preserved faecal samples after incubation in sM9 medium supplemented with 1965 μM entacapone for 48 hours under anaerobic conditions. An aliquot of the glycerol stock was serially diluted in PBS, and 10^-4^ and 10^-5^ dilutions were plated in BHIs agar. Isolated colonies were re-streak 3 times to purity and submitted to colony PCR using the 16S rRNA gene primers 616V (5’-AGA GTT TGA TYM TGG CTC AG-3’) and 1492R (5’-GGT TAC CTT GTT ACG ACT T −3’). Single colonies were picked with an inoculation loop, resuspended in 50 μl of nuclease free water and boiled at 95°C for 10 minutes to lyse the cells and release cell contents. After a short spin to pellet cell debris, 2 μl of the supernatant was added to a PCR reaction mix (final concentrations; Green 1X Dream Taq Buffer, dNTPs 0.2 mM, BSA 0.2 mg/ml, Taq polymerase 0.05 U/μl, primers 1 μM) prepared in a final volume of 50 μl per reaction. The amplification cycles were as follow: initial denaturation at 95 °C for 3 min, followed by 30 cycles at 95 °C for 30 s, 52 °C for 30 s, 72 °C for 1.5 min, and a final elongation at 72 °C for 10 min. PCR products were visualized on 1.5% agarose gel electrophoresis, and subsequently cleaned and concentrated on columns (innuPREP PCRpure Kit, Analytik Jena) according to the manufactureŕs instructions. Concentrations were measured by Nanodrop, and samples were sent for Sanger sequencing at Microsynth AG (Vienna, Austria). Sequencing results were analyzed by BLASTn^32^ against the 16S rRNA gene sequences retrieved by metagenomics. The near full 16S rRNA gene sequences of isolate E2 obtained were 99% identical to the 16S rRNA copies of *E. coli* D bin.187 and 100% identical to ASV_9ec_9n4.

### Growth of microbial pure cultures in the presence of entacapone and/or iron

*Bacteroides thetaiotaomicron* (DSM 2079) cells were grown anaerobically (85% N2, 10% CO2, 5% H2) at 37°C in *Bacteroides* minimal medium (BMM) containing 27.5 μM of iron (FeSO4, Merck)^91^. To test the effect of iron on rescuing the inhibitory effect of entacapone on *B. thetaiotaomicron* growth, entacapone-treated (1965 μM) cultures were supplemented with 100 μM, 500 μM or 1mM of iron. Growth rescue was observed when cultures were supplemented with either 500 μM FeSO4 (Fe(II)) or FeCl3 (Fe(III)), and for all subsequent experiments only FeSO4 was used. Cu(II)SO4 (500 μM or 1mM - Merck) or entacapone pre-complexed with iron (1965 μM, prepared as described above) were also supplemented to *B. thetaiotaomicron* cultures. Samples for total cell counts were taken at 0, 24, and 48 hours of growth under anaerobic conditions. *Phocaeicola dorei* (DSM 17855) was grown in BMM in the presence or absence of 1965 μM entacapone. After 24 and 48 hours of growth, an aliquot was collected and fixed with paraformaldehyde as described above, and stored at −20°C in PBS:EtOH 50% v/v until further analyses. To test if growth of the *Escherichia coli* faecal isolate E2 was dependent on the presence of iron, the isolate was grown in MOPS medium without or without 10 μM of FeSO4 in the presence or absence of entacapone for 24 hours at 37°C. To avoid the transfer of any iron from the pre-inoculum, pre-inoculum cells were washed thoroughly in MOPS medium without iron, prior to inoculation.

### Determination of total cell loads in pure cultures incubated with entacapone and iron

To determine microbial cell loads in pure culture, samples were collected at 0, 24 or 48 hours of growth and stained (either undiluted or after a dilution of 10 times in 1x PBS) with the QUANTOM™ Total Cell Staining Kit (Logos Biosystems, Korea). Total cell counts were determined using a QUANTOM Tx™ Microbial Cell Counter (Logos Biosystems, Korea), according to the manufacturer’s instructions.

## Data availability

The 16S rRNA gene sequences data have been deposited in the National Center for Biotechnology Information (NCBI) Sequence Read Archive (accession number PRJNA1033532).

## Code availability

CellProfiler pipelines and Matlab codes have been deposited in GitHub (https://github.com/buchenglab/srs-fish-drugs).

## Acknowledgements

Research reported in this manuscript was funded by NIH Awards R35GM136223 (to J.-X.C.) and R01AI141439 (to J.-X.C.) and supported by the Boston University Micro and Nano Imaging Facility and Office of the Director, NIH Award S10OD024993. The content is solely the responsibility of the authors and does not necessarily represent the official views of the NIH. Funding for the presented research was also provided via the Young Independent Research Group Grant ZK-57 (to F.C.P.) of the Austrian Science Fund (FWF) and the FWF-Wittgensteinaward Z383-B (to M.W.) as well as the FWF-funded Cluster of Excellence “Microbiomes drive Planetary Health” (COE 7). We thank Jasmin Schwarz, Gudrun Kohl, Petra Pjevac and Joana Seneca Silva from the Joint Microbiome Facility of the Medical University of Vienna and the University of Vienna for assisting with amplicon and metagenomic sequencing, as well as repositing of sequencing data.

## Author contributions

F.C.P., X.G., J.-X.C. and M.W. designed this study. F.C.P, X.G., J.M.K., K.M. and M.D. performed the experiments with input from T.B, Y.Z., D.B. and A.S.. F.C.P, J.M.K., R.H.K, B.H., and K.W. analyzed sequencing data and performed bioinformatic analysis. F.C.P. and X.G. wrote the manuscript with input from all co-authors and all authors read and approved the final manuscript.

## Ethics declarations

The authors declare no competing interests.

## Notes

### Competing Interest Statement

The authors have declared no competing interest.

### Summary of Updates

Authors affiliation updated

