## Supplementary Text and Figures for "The Parkinson’s drug entacapone disrupts gut microbiome homeostasis via iron sequestration"

**Included in this file:**

Supporting Information Text

Extended Data Figures 1 to 6

### Supporting Information Text

### SRS-FISH setup optimization

To further improve the SRS-FISH sensitivity and throughput, we have upgraded the platform in two major aspects. To increase sensitivity, we have switched from femtosecond SRS to picosecond SRS by spectral focusing[1,2] (Extended Data Fig.2; see Methods). Compared with femtosecond SRS, picosecond SRS allows the use of a higher laser power without damaging microbiota cells (Extended Data Fig.3a). Picosecond SRS also improves the ratio between the vibrational signal and the cross-phase modulation background when measuring Raman vibrational peaks with relatively narrow bandwidth. This enhancement is crucial for a more precise quantification of C-D bonds in metabolically active cells, that are formed through biotic incorporation of deuterium from heavy water (Extended Data Fig.3). Moreover, to increase the throughput when screening bacteria tagged by FISH, we combined point scanning SRS with widefield fluorescence imaging, which improved the screening efficiency, especially for rare cells.

### Chemical Imaging of Entacapone

To understand the non-vibrational resonant signal that we observed in the ENT-Hi treated samples, we first evaluated the optical properties of pure entacapone. Entacapone is an orange coloured, non-fluorescent compound exhibiting an absorption peak at 397 nm (Extended Data Fig.4a). Under the laser illumination of the SRS microscope, there are four types of light-matter interactions that potentially exist. That includes: vibrationally-resonant SRS, non-resonant cross-phase modulation (XPM), as well as electronically-resonant transient absorption and multiphoton photothermal (PT) that could be attributed to the 397 nm absorption peak of entacapone (Extended Data Fig.4a). As SRS and XPM are instantaneous, the signal is only generated when the pump and probe pulses are temporally overlapped (Extended Data Fig.4b). Transient absorption is mediated by electronic excited state with a lifetime of picosecond to nanosecond level, thus when tuning the delay time between the pump and probe pulses, the signal decays with a time constant on such a time scale[3]. The decay time of a PT signal is usually on the microsecond level[4], much longer than SRS, XPM and transient absorption. When the measurement is carried out at >1 MHz frequency, such long decay time could shift the phase of the signal to the y channel of the lock-in amplifier[5,6] (Extended Data Fig.4b, c), while optical delay (<30ps) could not affect the signal.

By evaluating the signal from ENT-Hi cells, we observed that when changing the delay time between pump and probe pulses, the x channel output of the lock-in amplifier resembles the cross-correlation function between pump and probe pulses, which is characteristic for an instantaneous signal such as SRS and/or XPM, while an almost constant signal was observed in the y channel, which is consistent with the long decay time of a PT signal (Extended Data Fig. 4d). Taken together, these results confirm that PT is the dominating origin of the signal (Extended Data Fig.4d).

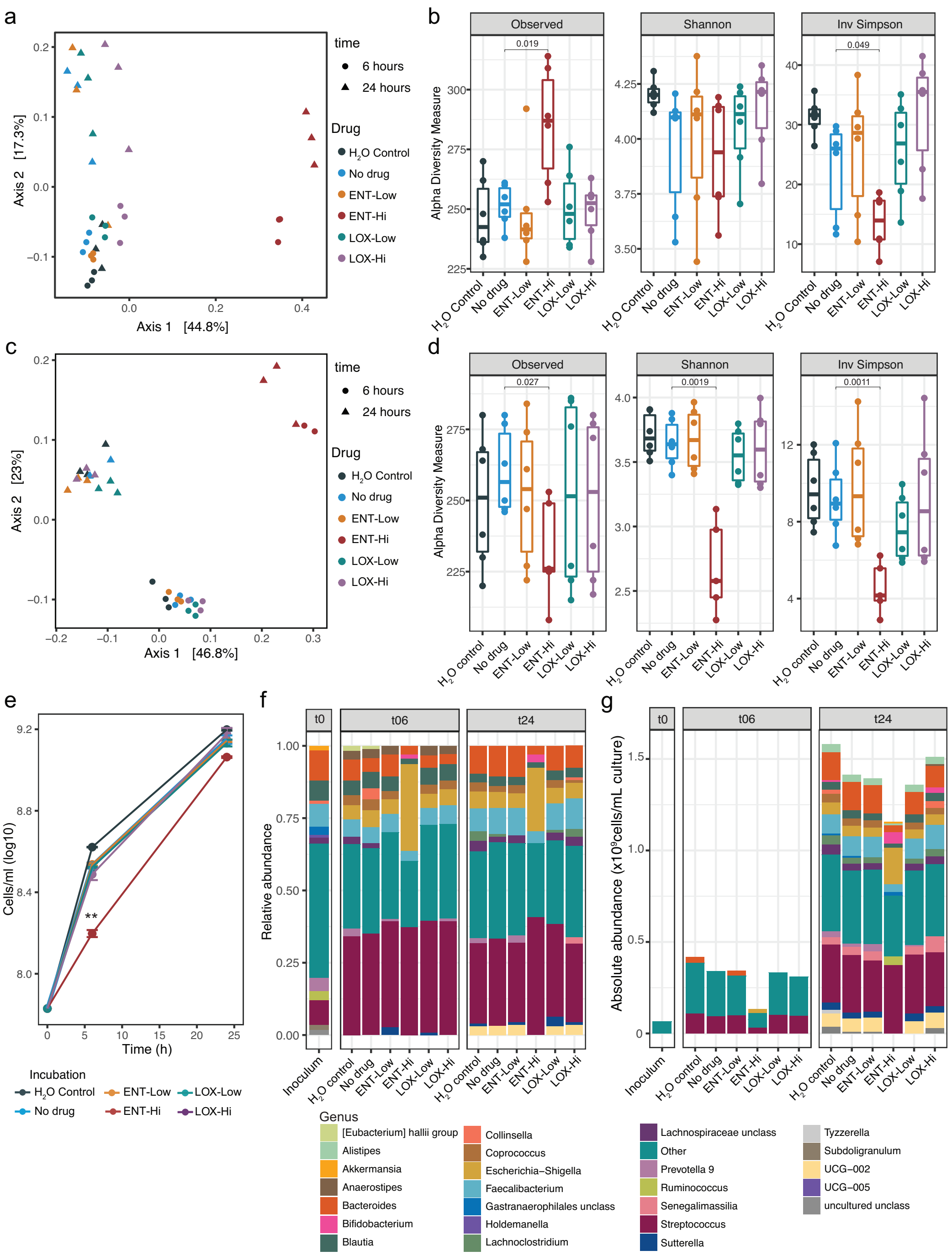


**Extended Data Fig.1. Relative and absolute abundance profiles of faecal samples resuspended in sM9 or BHI and incubated with drugs. a.** Ordination plot of Bray–Curtis distances of microbial communities (ASV-level composition) in faecal samples resuspended in sM9 medium (**a**) or BHI medium (**c**) and incubated in the presence of drugs. Samples are colored according to the drug amended and shaped according to the time of incubation. Please note that in (**a**) at T6 ENT-Hi condition two of the data points show overlapping positions in the ordination plot. sM9 medium: PERMANOVA =0.001, $R^2$=0.64. BHI medium: PERMANOVA =0.001, $R^2$=0.45. **b.** Alpha diversity metrics (number of Observed ASVs, Shannon Index, and Inverse Simpson Index) of faecal samples resuspended in sM9 medium (**b**) or BHI medium (**d**) and incubated in the presence of drugs. Each data point represents a replicate and samples from both 6 and 24 hours of incubation are shown. Significant differences determined by Wilcoxon testing are indicated. **e.** Total cell loads in faecal samples resuspended in BHI medium and incubated with drugs, as assessed by flow cytometry. **p<0.01; unpaired two-sample t-test with “No drug” used as a reference group. **f, g.** Relative (**f**) and absolute (**g**) genus abundance profiles of microbial communities originating from faecal samples resuspended in BHI medium and incubated in the presence of drugs at T0 and after 6 or 24 hours of incubation. All genera present at relative abundances below 0.025 or absolute abundances below $2.5x10^7$ cells.$ml^{-1}$ were assigned into the category “Other” (“unclass”: unclassified).

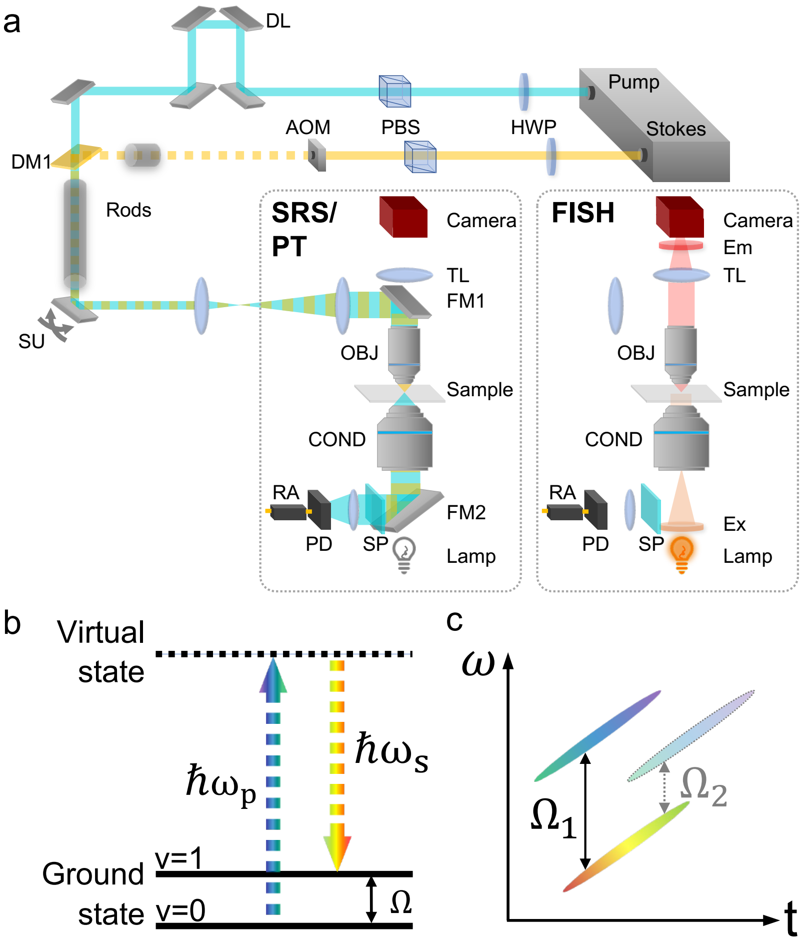


**Extended Data Fig.2. Pump probe microscopy combined with widefield fluorescence imaging for correlative chemical and fluorescence *in situ* hybridization imaging. a.** Experimental setup. Left dashed box: laser point scanning mode for SRS/PT. DL: delay line; HWP: half waveplate; PBS: polarized beam splitter; AOM: acousto-optic modulator; DM: dichroic mirror; SM: scanning unit; FM: flip mirror; OBJ: objective; COND: condenser; SP: short pass filter; PD: photodiode; RA: resonant amplifier; Ex: excitation filter; Em: emission filter; TL: tube lens. **b.** Stimulated Raman scattering energy diagram. $\omega_p$: pump beam frequency; $\omega_s$: Stokes beam frequency; $\hbar$: Planck's constant; $\Omega$: energy of the vibrational mode; v: vibrational level. **c.** Picosecond stimulated Raman scattering by spectral focusing.

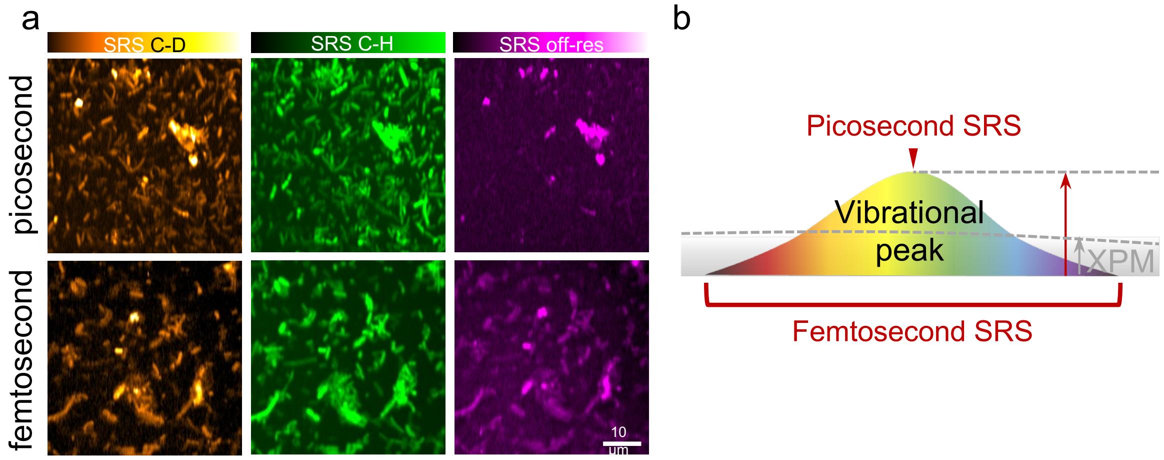


**Extended Data Fig.3. Sensitive mapping of microbial activity by picosecond SRS. a.** Picosecond SRS alleviates cross-phase modulation (XPM) and provides higher sensitivity for measuring the activity of microbial cells via deuterium incorporation with a laser power below the cell-damaging threshold. Picosecond SRS pump power: 30 mW, Stokes power: 120 mW. Femtosecond SRS: pump power: 15 mW, Stokes power: 70 mW. **b.** Picosecond and femtosecond SRS probing difference in the spectral domain. For picosecond SRS, the targeting bandwidth is 20 $cm^{-1}$, which is much narrower than the femtosecond SRS which covers more than 200 $cm^{-1}$. Thus, the ratio between the signal (vibrational feature) and the background (cross phase modulation, XPM) is improved in picosecond SRS.

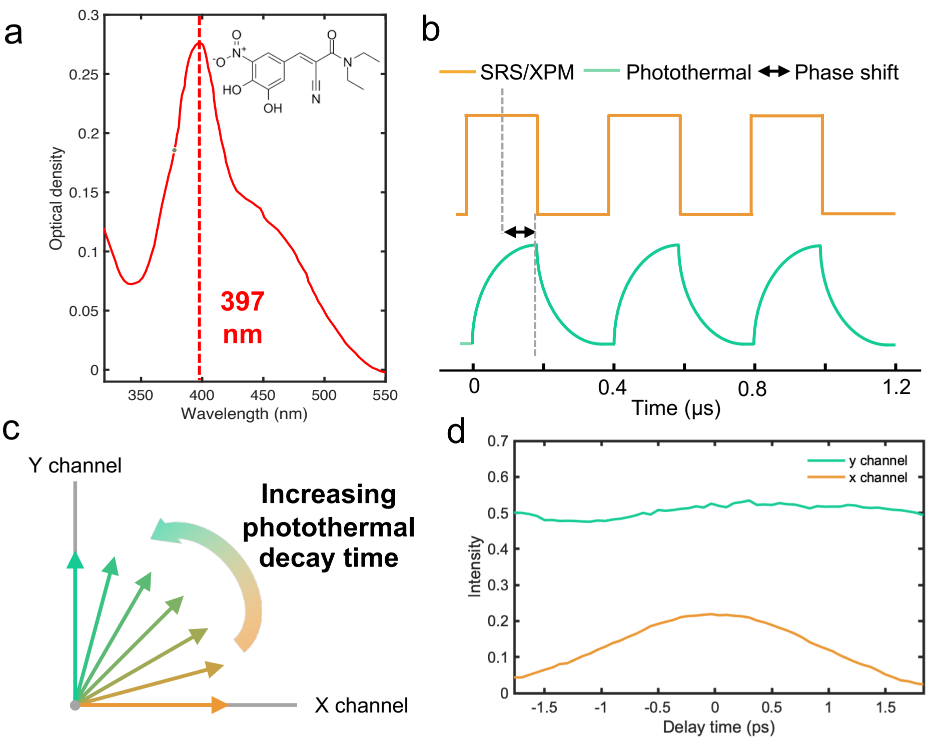


**Extended Data Fig.4. Entacapone photothermal signal retrieved from modulation transfer detection. a.** 10 mM Entacapone DMSO solution (under acidic pH conditions) measured by UV-VIS spectroscopy. **b**. Photothermal signal time trace exhibits a π/2 phase delay relative to the intensity modulation by the AOM and instantaneous signals, such as SRS and XPM. **c.** Pump probe signal phase upon increasing photothermal decay time. **d.** Entacapone accumulation in microbiota cells via pump probe detection with delay in time and x/y channel lock-in amplifier signal detection. As photothermal is not sensitive to picosecond level delay, the y channel showed a constant strong photothermal signal from the drug, while in the x-channel, the instantaneous cross phase modulation background showed up as a cross-correlation profile of two the pump and probe laser beams.

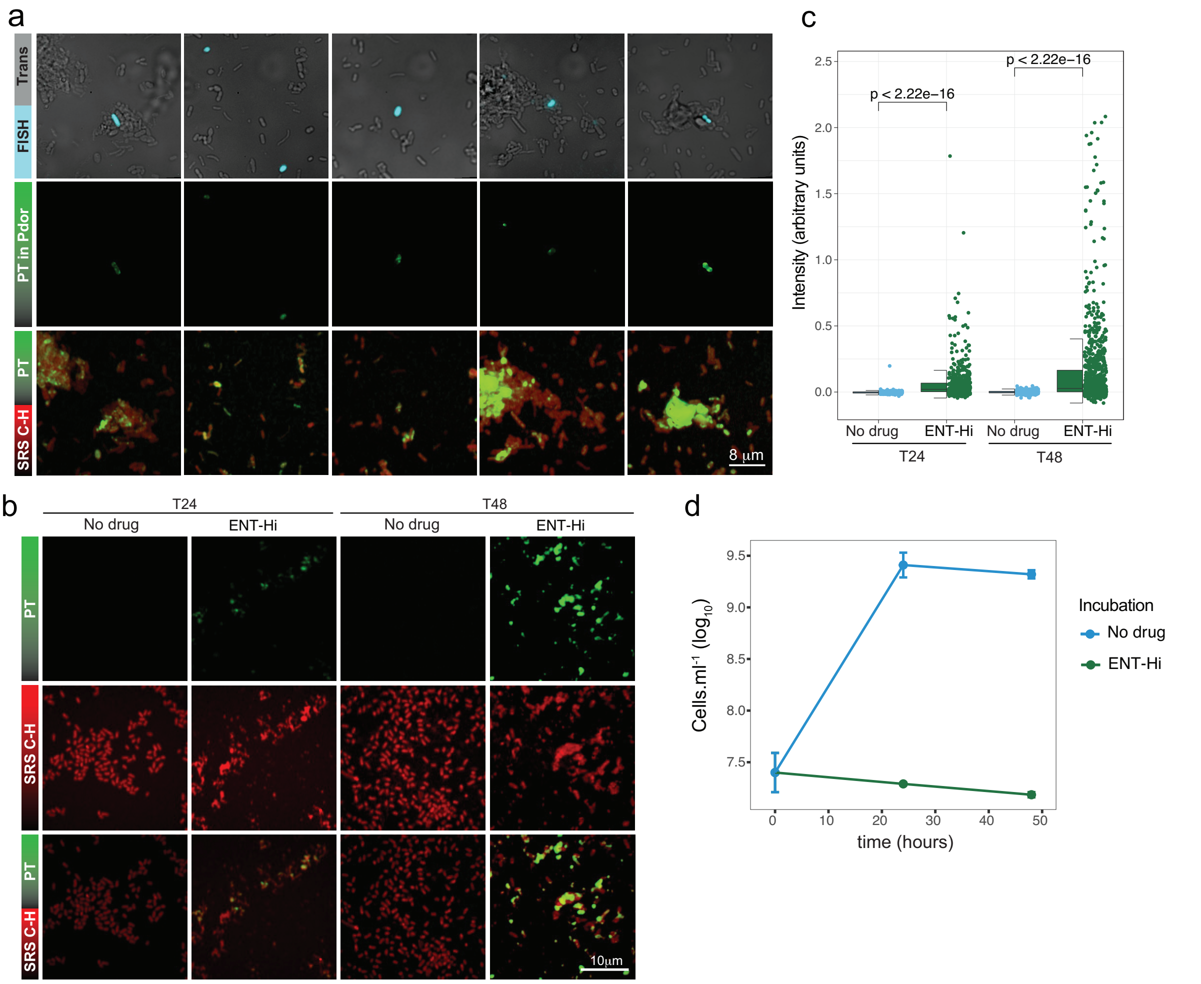


**Extended Data Fig.5. Photothermal imaging of entacapone accumulation by *Phocaeicola dorei*.** **a.** All columns show representative images of faecal samples incubated with ENT-Hi for 6 hours followed by hybridization with a fluorescently-labeled oligonucleotide probe targeting *Phocaeicola dorei* (Pdor) (see Supplementary Table 9). Probe signals from hybridized cells (displayed in cyan) were detected using widefield fluorescence microscopy (top panel). Photothermal signal intensity from entacapone (PT, displayed in green) in Pdor cells only (PT signals of other cells were removed to enhance clarity) is shown in the middle panel. In the bottom panel all PT signals are displayed and overlayed with the SRS C-H signals (displayed in red). PT channel contrast: min 0 max 1.8. C-H channel is represented on a log scale. **b.** Representative images of the pure culture *Phocaeicola dorei* strain 175 incubated anaerobically in growth medium with entacapone for 24 or 48 hours. Photothermal signal from entacapone (PT, displayed in green) in *P. dorei* cells is shown in the top panel, SRS C-H signal (displayed in red) in the middle panel, and an overlay of both signals is shown in the bottom panel. **c.** Single-cell specific photothermal signal intensity detected in samples shown in b. Boxes represent the median, first and third quartile. Whiskers extend to the highest and lowest values that are within one and a half times the interquartile range. p-values were determined by the unpaired two-sample t-test. **d.** Growth (assessed by microbial cell counts) of *P. dorei* 175 in the presence or absence of entacapone.

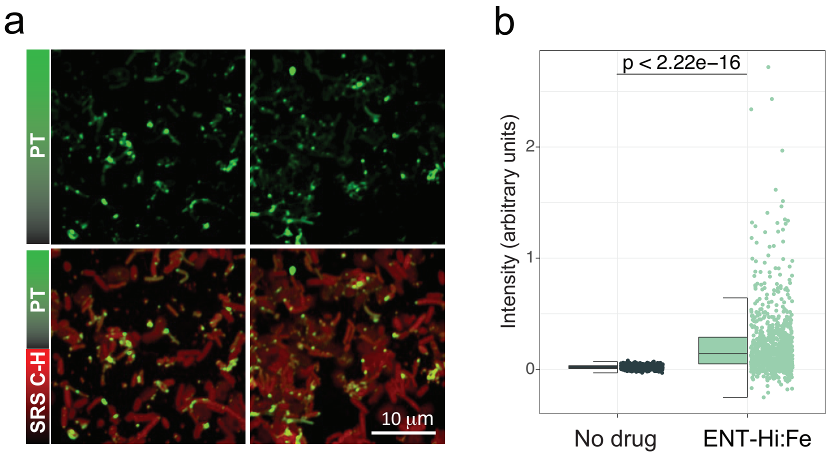


**Extended Data Fig.6. Accumulation of entacapone pre-complexed with Fe(II) by microbiome cells. a.** Both columns show representative images of faecal samples anaerobically incubated for 6 hours with ENT-Hi pre-complexed with 1 mM $FeSO_4$ (ENT-Hi:Fe). Photothermal signal from entacapone (PT, displayed in green) is shown on top and the overlay with the SRS C-H biomass signal (displayed in red) is shown in the bottom. **b.** Single-cell specific photothermal signal intensity in samples shown in b. Boxes represent median, first, and third quartile. Whiskers extend to the highest and lowest values that are within one and a half times the interquartile range. p-value was determined with the unpaired two-sample Wilcoxon test.
